# The extra-terminal domain drives the role of BET proteins in transcription

**DOI:** 10.1101/2025.05.12.653540

**Authors:** Iwona Pasionek, John B. Ridenour, Agnieszka Machowska, Magdalena Donczew, Michael T. Kinter, Kevin A. Boyd, Rafal Donczew

## Abstract

BET proteins facilitate the transcription of most eukaryotic genes, yet the specific mechanisms underlying their function remain incompletely understood. As chromatin readers, BET proteins use tandem bromodomains to recognize acetylated lysine residues on histones and other protein partners. However, recent evidence indicates that bromodomain activity alone does not account for the full spectrum of BET protein functions, underscoring the importance of additional conserved domains. Here, we systematically evaluated all conserved domains of BET proteins and identified the extra-terminal (ET) domain as essential for cell viability, genome-wide transcription, and BET chromatin occupancy. Moreover, we demonstrate that the ET domain exerts these effects by acting as a central hub for interactions with multiple transcriptional regulators. Our findings advance the current understanding of BET protein biology and reveal potential mechanisms by which cells can evade bromodomain inhibition under pathological conditions.

## Introduction

Bromodomains (BDs) are reader modules that recognize acetylated lysine residues on histones and other protein targets^1^. Members of the bromodomain and extra-terminal domain (BET) family are conserved chromatin readers that play central roles in transcription regulation^2–5^. Consistent with this function, BET proteins have been implicated in sustaining pathological transcriptional programs across various human diseases^6,7^. Pharmacological inhibition of BET bromodomains represents a promising therapeutic strategy for cancer and other disorders; however, resistance to BD inhibitors frequently emerges, highlighting the need for alternative approaches to target BET proteins^8–12^. However, a comprehensive model describing how BET proteins regulate transcription remains elusive.

All three mammalian BET proteins expressed in somatic tissues (BRD2, BRD3, and BRD4) contribute to gene transcription, but BRD4 functions as the dominant BET factor, with primary roles in transcription elongation^3–5,13,14^. Budding yeast encode two BET proteins (Bdf1 and Bdf2, referred to here as Bdf1/2), at least one of which is required for viability. Bdf1/2 make overlapping contributions to transcription; however, similar to BRD4 in mammalian cells, yeast cells rely predominantly on Bdf1^2,15^. Mechanistically, Bdf1/2 support both the initiation and elongation phases of transcription^2^. Notably, mammalian BET proteins have also been implicated in transcription initiation, indicating that BET-dependent transcriptional functions are broadly conserved across eukaryotic lineages^4,14,16^.

According to the canonical model, the tandem BDs serve as the primary module supporting BET protein function^1,8,17^. The multiply acetylated N-terminal tail of histone H4 is a preferred target of most BET BDs, including both BDs of Bdf1^14,18,19^. However, we previously showed that rapid depletion of H4 acetylation results in only an ∼50% reduction in Bdf1 chromatin occupancy^2^. Consistent with this observation, yeast cells remain viable in the absence of acetylated residues on the H4 tail^20^; Bdf1 is recruited to immobilized promoter DNA even in the absence of nucleosomes^15,21^; a Bdf1 mutant with inactivated BDs suppresses the temperature sensitivity caused by *BDF1* deletion^18^; and chemical inhibition of BET BDs in mammalian cells neither fully displaces BET proteins from chromatin nor abolishes their function^4,5,16,22^. Together, these findings indicate that BET-mediated transcriptional regulation involves additional, yet poorly understood mechanisms that do not depend on BD activity.

BET proteins exhibit a modular architecture composed of conserved structured regions (BDs, motif B, and the extra-terminal (ET) domain) and conserved unstructured regions [the N- and C-terminal phosphorylation sites (NPS and CPS) and the basic interaction domain (BID)], interspersed with additional unstructured regions lacking sequence conservation (**Fig. 1A**). Although the functions of non-BD domains have not been comprehensively examined, existing evidence suggests that they play important roles. For instance, the ET domain of mammalian BET proteins serves as an interaction hub for multiple protein partners^17,23–25^, and these interactions have been proposed to contribute to BRD4-mediated transcriptional regulation, BRD4 chromatin recruitment, and the survival of certain cancer cell lines^22,23,26–29^. However, the *in vivo* significance of the ET domain for BET protein function remains unresolved. In addition, phosphorylation within the NPS and CPS regions has been implicated in enhancing BET association with chromatin and protein partners^10,11,30^. Finally, motif B and the BID have been biochemically shown to promote BET protein dimerization^31^.

**Fig. 1.**
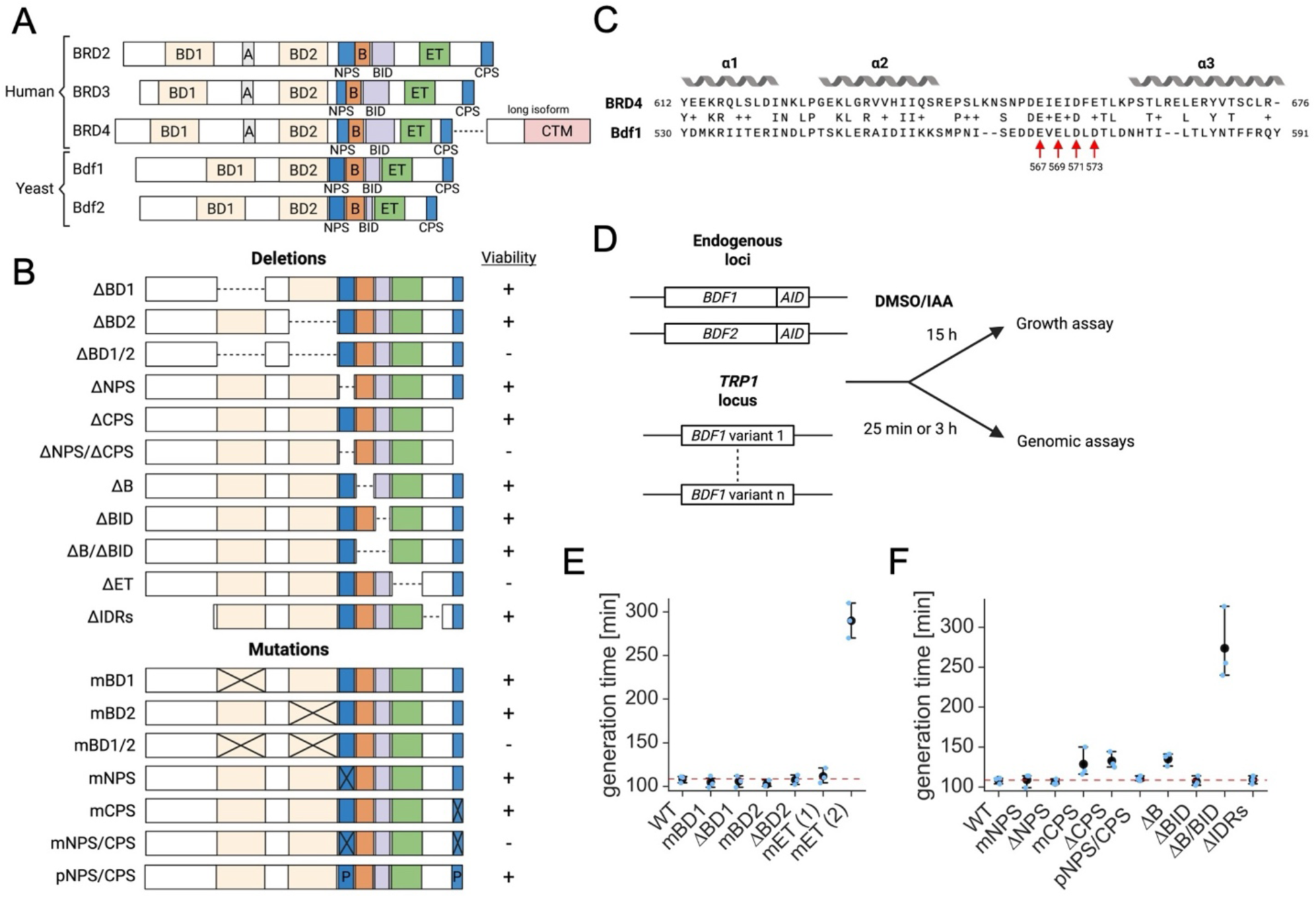
The BET protein ET domain is essential for viability. (**A**) Conserved domain organization of BET proteins. BD – bromodomain, A/B – motifs A/B, BID – basic interaction domain, ET – extra-terminal domain, NPS/CPS – N/C-terminal phosphorylation sites, CTM – C-terminal motif (BRD4 long isoform only). (**B**) *BDF1* variants constructed in this work. Ability to support growth in the absence of endogenous *BDF1/2* is indicated. (**C**) Sequence alignment of the ET domain of BRD4 (top) and Bdf1 (bottom). Four conserved acidic residues important for interactions between BRD4 and protein partners are marked. Bdf1 coordinates are shown below the arrows. (**D**) Schematic representation of the experimental strategy for viability and genomic (SLAM-seq and ChEC-seq) assays. Experiments were performed in strains in which endogenous Bdf1/2 carried auxin-inducible degron (AID) tags and unmodified *BDF1* or *BDF1* variants were expressed from the *TRP1* locus. Endogenous Bdf1/2 were depleted by the addition of 3-indoleacetic acid (IAA). (**E-F**) Growth assays comparing the fitness of a control strain expressing unmodified *BDF1* (WT) to strains expressing *BDF1* variants after 15 h exposure to IAA (long-term loss of endogenous Bdf1/2). Black markers indicate mean generation time. Error bars represent 95% confidence intervals (*n* = 3).

Here, we delineated the contributions of the conserved domains of BET proteins to cell viability, transcription, BET chromatin association, and the recruitment and/or maintenance of partner proteins. Our key findings establish the ET domain as essential for BET protein function in living cells and enable us to propose a revised model of BET-mediated transcriptional regulation.

## Results

### The BET protein ET domain is essential for cell survival

Prior yeast genetic studies established that the BDs and NPS/CPS regions of BET proteins contribute to cell viability, but the functional significance of the remaining conserved domains has not been defined^18,32^. We previously demonstrated that although Bdf1 and Bdf2 have redundant roles in transcription, Bdf1 functions as the dominant BET protein in yeast^2^. Accordingly, we selected Bdf1 as our model BET factor. We generated a comprehensive collection of *BDF1* variants containing deletions or targeted mutations across all conserved BET domains. When previous work suggested functional overlap, we included combinatorial perturbations affecting more than one domain. Because intrinsically disordered regions (IDRs) have been implicated in BRD4 function in mammalian cells, we also constructed a *BDF1* variant with truncations in IDRs that do not overlap with the conserved unstructured NPS, CPS, or BID regions^33^. For the BDs and NPS/CPS regions, we introduced targeted mutations in residues required for BD function or in serine residues with documented phosphorylation, respectively^18,32^. As a comparison, we also generated a phosphomimetic *BDF1* variant in which all phosphorylated serines were substituted with glutamate. In total, we constructed eighteen *BDF1* variants (**Fig. 1B**). To facilitate stable expression in yeast cells, we cloned *BDF1* variants or unmodified *BDF1* into a minichromosomal plasmid pRS315 under the control of endogenous *BDF1* promoter and 3’ untranslated region (**Table S1**).

We performed a complementation assay in a strain dependent solely on *BDF1* expressed from a minichromosomal plasmid, as endogenous *BDF1/2* were deleted. To test whether each *BDF1* variant could support viability, we attempted to replace the plasmid carrying unmodified *BDF1* and the URA3 marker with plasmids encoding the *BDF1* variants, selecting for viable transformants on medium containing 5-fluoroorotic acid (5-FOA), which counterselects URA3-containing cells. Five *BDF1* variants failed to support viability: the double BD-deletion and BD-mutation variants, the variants lacking both NPS and CPS regions or carrying serine-to-alanine substitutions across both regions, and the variant lacking the ET domain (**Fig. 1B**). Proper expression of these *BDF1* variants was validated in experiments described below. In summary, although prior studies established *in vivo* requirements for the BDs and NPS/CPS regions of BET proteins in yeast^18,32^, our findings reveal for the first time that the ET domain is essential for cell viability.

### A conserved stretch of acidic residues in the ET domain is required for its function

A conserved cluster of acidic residues located between the second and third helices of the ET domain has been shown to mediate BRD4 interactions with several protein partners^23–25^. To assess the *in vivo* significance of this region in Bdf1, we generated six additional *BDF1* variants in which different numbers of four conserved acidic residues (amino acids 567, 569, 571, and 573) were substituted with alanine (**Figs. 1C and S1A**). Using the complementation assay described above, we found that mutation of three or more of these acidic residues is lethal in yeast, demonstrating that this region of the ET domain is essential for its function. For subsequent experiments, we selected a representative set of ET-domain mutants consisting of a single substitution (D571A; mET(1)), a double substitution (D571A, D573A; mET(2)), and a quadruple substitution (mET(4)) (**Figs. 1C and S1A**).

To support subsequent experiments, we integrated all *BDF1* variants, along with an unmodified *BDF1* control, into the TRP1 locus of a strain in which endogenous Bdf1/2 carry auxin-inducible degron (AID) tags (RDY73; Bdf1/2-AID strain) (**Figs. 1D, and Tables S1 and S2**)^2^. In this system, each *BDF1* variant is constitutively expressed from an ectopic locus under the control of the endogenous *BDF1* promoter and 3′ untranslated region, while endogenous Bdf1/2 can be rapidly depleted upon addition of 3-indoleacetic acid (IAA)^2^. Importantly, integration of an additional unmodified *BDF1* copy did not affect cell viability, and the *BDF1* variants were expressed at endogenous or modestly elevated levels, both in the presence and absence of endogenous Bdf1/2 (**Fig. S1B-D**). We did not detect expression of variants containing the BD2 deletion (ΔBD2, ΔBD1/2) or the deletion of the N-terminal unstructured region (ΔIDRs), suggesting that the antibody used recognizes epitopes located within these regions of Bdf1. Nevertheless, expression of the ΔBD2, ΔBD1/2, and ΔIDRs variants is supported by normal growth rates (ΔBD2 and ΔIDRs variants) and by the modest transcriptional changes observed in cells that rely on these *BDF1* variants, as described below.

The complementation assay identified the BET protein domains required for viability. To extend these findings, we quantified growth-rate defects associated with each *BDF1* mutation. Yeast cells were cultivated in the presence of IAA, forcing dependence on the Bdf1 variants as endogenous Bdf1/2 were depleted, and generation times were calculated after ∼15 hours of growth (**Fig. 1D**). First, strains harboring *BDF1* variants that failed to support viability in the complementation assay displayed complete growth inhibition, validating our earlier results. Second, whereas a single point mutation in the ET domain was well tolerated, mutation of only two conserved acidic residues caused a severe growth defect. By contrast, deletion or mutation of a single BD did not impair growth (**Fig. 1E**). Third, we found that CPS, but not NPS, deletion or mutation modestly slowed growth. Notably, the NPS/CPS phosphomimetic Bdf1 variant fully supported viability, suggesting that BET proteins function optimally in a fully phosphorylated state and that their phosphorylation may not be dynamically regulated under normal growth conditions (**Fig. 1F**). Fourth, combined loss of the B/BID regions substantially impaired growth, consistent with their proposed role in promoting BET dimerization^31^. Finally, truncation of the Bdf1 IDRs did not affect viability, indicating that the large, unstructured N-terminal region of BET proteins does not play a major biological role (**Fig. 1F**). As an additional validation, we observed similar growth rates across all viable strains relying solely on *BDF1* variants expressed from minichromosomal plasmids (**Fig. S1E and Table S2**). Collectively, these assays define the contributions of all conserved BET protein domains to yeast viability and identify a critical functional region within the ET domain.

### The ET domain is essential for genome-wide transcription

The essential function of BET proteins is to sustain global transcriptional programs^2–5^. To define the contribution of each conserved BET domain to genome-wide transcription, we analyzed our collection of strains expressing *BDF1* variants in the background of endogenous Bdf1/2 fused to AID tags (**Fig. 1D and Table S2**). We previously confirmed that introducing an additional unmodified copy of *BDF1* does not compromise viability and that Bdf1 protein levels are comparable when expressed from an ectopic versus endogenous locus (**Fig. S1B-D**). We further validated that this additional *BDF1* copy does not substantially alter baseline transcription and largely rescues the transcriptional defects caused by depletion of endogenous Bdf1/2 (**Figs. S2A-B**). Finally, to assess whether any *BDF1* variants exert dominant-negative effects in the presence of endogenous Bdf1/2, we examined growth phenotypes and found that the two strains harboring variants with combined deletion or mutation of both NPS/CPS regions displayed pronounced slow-growth phenotypes (**Fig. S2C**).

In contrast to the growth assay, where cells had to rely on mutant Bdf1 variants for several generations, here we depleted endogenous Bdf1/2 for 25 minutes, followed immediately by 4-minute labeling of newly synthesized RNA with 4-thiouracil to facilitate quantification using SLAM-seq^34^ (**Fig. 1D**). This approach allowed us to track direct consequences of Bdf1 mutations, including lethal mutations, with minimized bias from indirect effects that accumulate during extended exposure to adverse genetic variants. We performed three replicate experiments for all strains with (IAA) or without (dimethyl sulfoxide (DMSO)) depletion of endogenous Bdf1/2 (**Table S3**). Variability in RNA levels between replicates in the final dataset (see Materials and Methods) was <30–40% for most genes (**Figs. S3A-B and Table S3**). We quantified transcriptional changes associated with perturbations in BET protein domains by comparing strains expressing *BDF1* variants to the strain expressing unmodified *BDF1* following depletion of endogenous Bdf1/2 (IAA treatment) (**Table S3**). To identify dominant-negative effects, we compared strains expressing *BDF1* variants without depletion of endogenous Bdf1/2 to the Bdf1/2-AID parental strain (DMSO treatment), which lacks an additional *BDF1* copy (**Fig. S3C and Table S3**). Consistent with the growth assay (**Fig. S2C**), strains expressing *BDF1* variants with mutations or deletions in both NPS/CPS regions (mNPS/CPS and ΔNPS/CPS) exhibited substantial transcriptional deviations (median change > 2-fold) in the presence of endogenous Bdf1/2. We observed a similar dominant-negative phenotype for the strain expressing the CPS deletion variant (ΔCPS). Consequently, we excluded ΔCPS, mNPS/CPS and ΔNPS/CPS strains from further analysis and experiments.

We aligned the transcriptional changes measured following depletion of endogenous Bdf1/2 with those observed upon loss of Bdf1/2 in the Bdf1/2-AID strain. Surprisingly, only *BDF1* variants lacking the ET domain or carrying mutations in all four conserved acidic residues of the ET domain (hereafter referred to as the ET domain mutation) produced substantial transcriptional defects (median decreases of 3.6-fold and 2.6-fold, respectively). By contrast, deletion or mutation of both BDs resulted in relatively modest transcriptional changes (median decreases of 1.5-fold and 1.4-fold, respectively) (**Figs. 2A and S4A**). Notably, the transcriptional defects caused by loss of ET domain function were comparable to those resulting from loss of Bdf1/2 in the Bdf1/2-AID strain (median decrease of 3.7-fold). Analysis of transcriptional changes at the level of individual genes and across previously defined major gene classes further confirmed that loss of ET domain function closely phenocopies the loss of BET proteins^35^ (**Figs. 2B-C, S4B and Table S3)**. Perturbations in other domains produced only minor effects on genome-wide transcription, with the exception of the CPS mutation and the combined deletion of the B/BID regions, both of which caused moderate decreases (median decreases of 1.7-fold and 1.6-fold, respectively) (**Fig. S4A**).

**Fig. 2.**
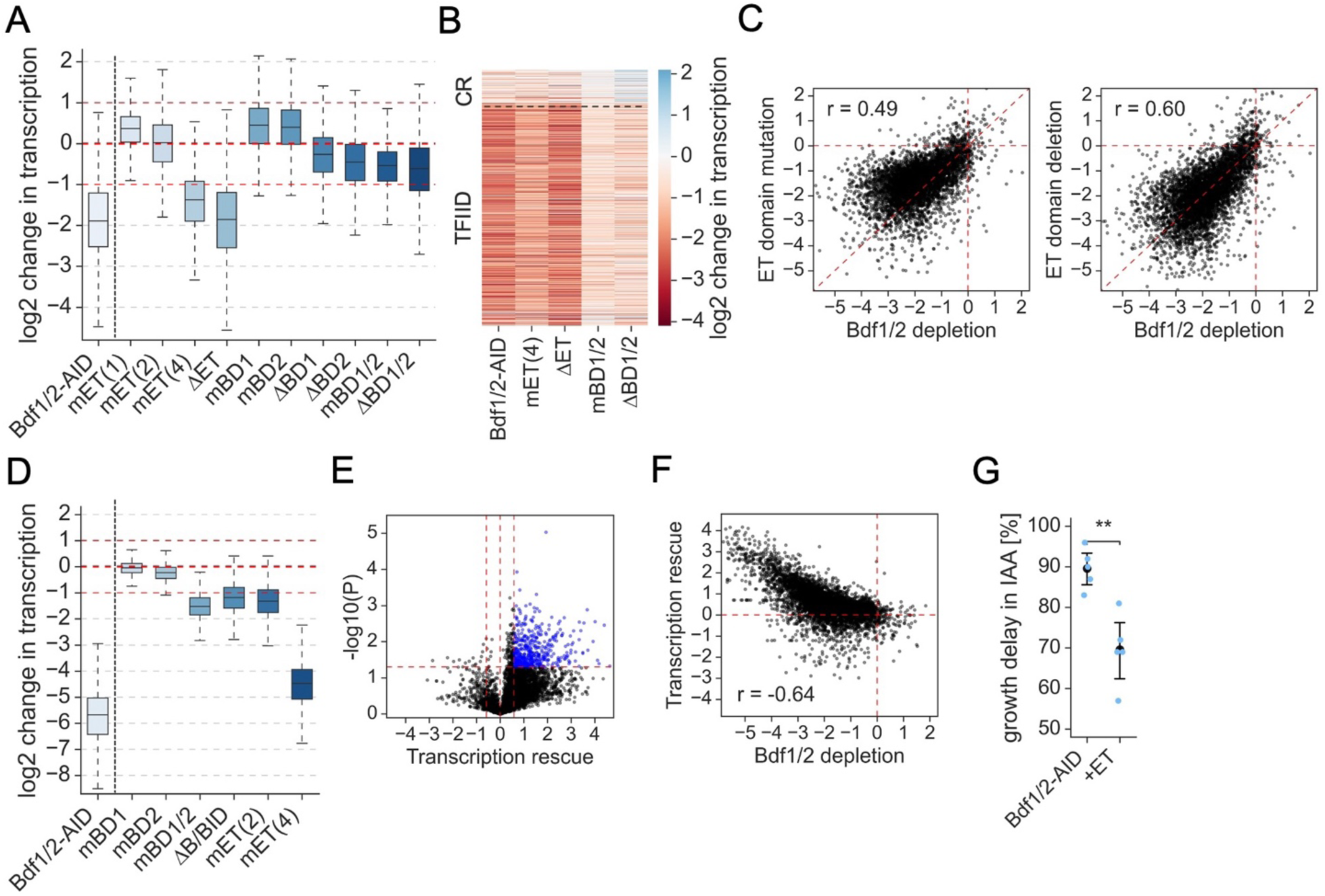
The ET domain is required for transcription of most genes. (**A**) Log2 changes in transcription due to selected *BDF1* mutations measured by SLAM-seq after depleting endogenous Bdf1/2 for 25 min (*n* = 4,836). Results of depletion of Bdf1/2 in the absence of additional copy of *BDF1* are shown in the first sample from the left. Results for additional *BDF1* variants are shown in **Fig. S4A**. (**B**) Heatmap representation of selected results from (**A**). Genes are divided into previously defined major yeast gene classes (CR – coactivator-redundant, TFIID – TFIID-dependent)^35^ (*n* = 4,560). (**C**) Comparison of selected results from (**A**). Pearson correlation coefficient (r) is shown (*n* = 4,836). (**D**) Log2 changes in transcription due to *BDF1* mutations after depleting endogenous Bdf1/2 for 3 h (*n* = 4,457). Results of depletion of Bdf1/2 in the absence of additional copy of *BDF1* are shown in the first sample from the left. (**E**) Volcano plot comparing the ability of the ET domain to rescue transcriptional changes following Bdf1/2 loss (log2 fold change over control strain) with associated *p*-values (*n* = 4,715). Thresholds used: fold change – 1.5-fold; *p*-value – 0.05 (Welch’s t-test). (**F**) Comparison of ET domain-mediated rescue of transcription following Bdf1/2 loss (y-axis, log2 fold change over control strain) with gene dependence on Bdf1/2 (x-axis, log2 scale). Pearson correlation coefficient (r) is shown (*n* = 4,715). (**G**) Growth assay comparing the fitness of the Bdf1/2-AID strain and the ET domain overexpressing strain during endogenous Bdf1/2 depletion. Samples were taken 1.5 and 3 h after DMSO or IAA addition, and growth delay was calculated by comparing generation times of IAA-treated and DMSO-treated cells. Black markers indicate mean growth delay. Error bars represent 95% confidence interval (*n* = 5). Results of a Welch’s t-test are shown (** – *p*-value < 0.01).

A modest change in transcription following BD inactivation was unexpected given the lethal phenotype associated with this mutation. We observed a similar discrepancy for deletion of the B/BID regions and mutation of two conserved acidic residues in the ET domain, both of which produced severe growth defects yet only minor transcriptional changes. To validate these findings, we extended the depletion period of endogenous Bdf1/2 to 3 hours to assess the accumulation of indirect effects on transcription. After 3 hours of IAA treatment, the Bdf1/2-AID strain exhibited a near-complete loss of genome-wide transcription (median decrease of 51-fold) (**Fig. 2D and Table S3**). In contrast, BD1/2 mutation, ET domain double mutation, and B/BID deletion resulted in more modest median decreases of 2.9-fold, 2.5-fold, and 2.3-fold, respectively. Compared to the rapid depletion experiment (**Figs. 2A and S4A**), these results indicate that the viability defects associated with BD1/2 mutation, the ET domain double mutation, or B/BID deletion arise primarily from indirect transcriptional dysfunction. Nevertheless, the indirect effects observed after 3 hours of IAA treatment were substantially stronger for the ET domain quadruple mutation (median decrease of 22.3-fold), consistent with trends observed following 25-minute depletion of endogenous Bdf1/2. In comparison, mutation of individual bromodomains did not appreciably affect transcription after prolonged IAA treatment, consistent with the minimal viability defects associated with these *BDF1* variants.

Based on our findings, we hypothesized that the ET domain can at least partially compensate for the loss of BET proteins. To test this hypothesis, we overexpressed the ET domain in the Bdf1/2-AID strain (**Fig. S4C**) and quantified transcriptional changes following 25 minutes of induced Bdf1/2 depletion. First, we observed that very few genes exhibited substantial transcriptional changes as a result of ET domain overexpression in the presence of endogenous Bdf1/2 (**Fig. S4D-E and Table S3**). Next, we compared transcriptional responses to Bdf1/2 loss between the ET domain overexpressing strain and the parental Bdf1/2-AID strain. We found that the ET domain compensated for the loss of BET proteins at many genes, with 455 genes showing statistically significant transcriptional rescue (**Fig. 2E and Table S3**). Notably, this effect was most pronounced at strongly Bdf1/2-dependent genes (**Fig. 2F**). Consistent with these observations, the ET domain partially rescued the growth phenotype associated with BET protein loss, although prolonged exposure to IAA ultimately remained lethal (**Fig. 2G**). In summary, these results reveal a surprisingly small direct contribution of the BDs to transcription and demonstrate that the ET domain is a dominant BET protein domain in transcriptional regulation.

### The ET domain facilitates BET protein recruitment to chromatin

To modulate transcription, BET proteins first need to be recruited to gene regulatory elements. Experiments in both yeast and mammalian cells have shown that BD function does not fully account for BET protein chromatin occupancy^2,4,15,16,22^. To identify BET protein domains that contribute to Bdf1 recruitment to chromatin, we modified selected yeast strains used in the SLAM-seq experiments by inserting a micrococcal nuclease (MNase) coding sequence before the stop codon of *BDF1* variants to facilitate ChEC-seq analysis (**Tables S1 and S2**)^36^. We previously found that ChEC-seq complements and outperforms ChIP-seq for mapping BET protein chromatin occupancy in yeast^2^. First, we validated that ectopically expressed Bdf1-MNase supports normal growth in the presence of AID-tagged endogenous *BDF1*/2 (**Fig. S5A**). We also confirmed that ectopically expressed Bdf1-MNase displays a chromatin-binding pattern highly similar to that of Bdf1-MNase expressed from the endogenous locus, both in the presence and absence of endogenous Bdf1/2 (**Fig. S5B**). As in the SLAM-seq assays, sample collection was preceded by a 25-minute depletion of endogenous Bdf1/2 to minimize confounding effects from endogenous BET proteins. All experiments were performed in three biological replicates, and the replicate datasets were highly consistent (**Fig. S5C**).

To evaluate the impact of targeted mutations in BET protein domains on Bdf1 recruitment, we compared data collected from strains expressing *BDF1* variants to that from the control strain expressing unmodified *BDF1* in the context of endogenous Bdf1/2 depletion (IAA treatment) (**Table S4**). Consistent with the major impact of the ET domain on transcription, we found that this domain broadly contributed to *Bdf1* chromatin occupancy (**Figs. 3A-C**). Surprisingly, at most gene promoters, the contribution of the ET domain to Bdf1 occupancy exceeded that of the BDs. Notably, deletion of the BDs or the ET domain produced very similar effects (**Figs. 3D and S6A-C**). Among other domains, perturbations in the CPS domain and the B/BID regions also affected Bdf1 recruitment, indicating that BET protein phosphorylation and dimerization both contribute to establishing genome-wide BET occupancy (**Fig. S6D-F**). None of the examined mutations caused substantial redistribution of Bdf1 to new loci. Next, we compared changes in Bdf1 promoter recruitment resulting from mutation of the BDs or the ET domain with levels of histone H4 lysine 12 acetylation (H4K12ac), a preferred target of Bdf1 BDs^19^. As expected, changes in Bdf1 recruitment due to BD mutation showed a modest correlation with H4K12ac levels, consistent with prior findings that BD inhibitors preferentially evict BET proteins from loci with high H4 acetylation, such as super enhancers^4^. In contrast, the effects of the ET domain on Bdf1 occupancy were independent of promoter acetylation levels (**Fig. 3E**). Finally, we observed only a weak correlation between changes in Bdf1 recruitment and transcriptional output (**Fig. S6G**), consistent with the previously reported lack of correlation between baseline Bdf1 promoter occupancy and transcription^2^. In summary, we revealed that the ET domain, not the BDs, is the primary contributor to the BET protein chromatin occupancy.

**Fig. 3.**
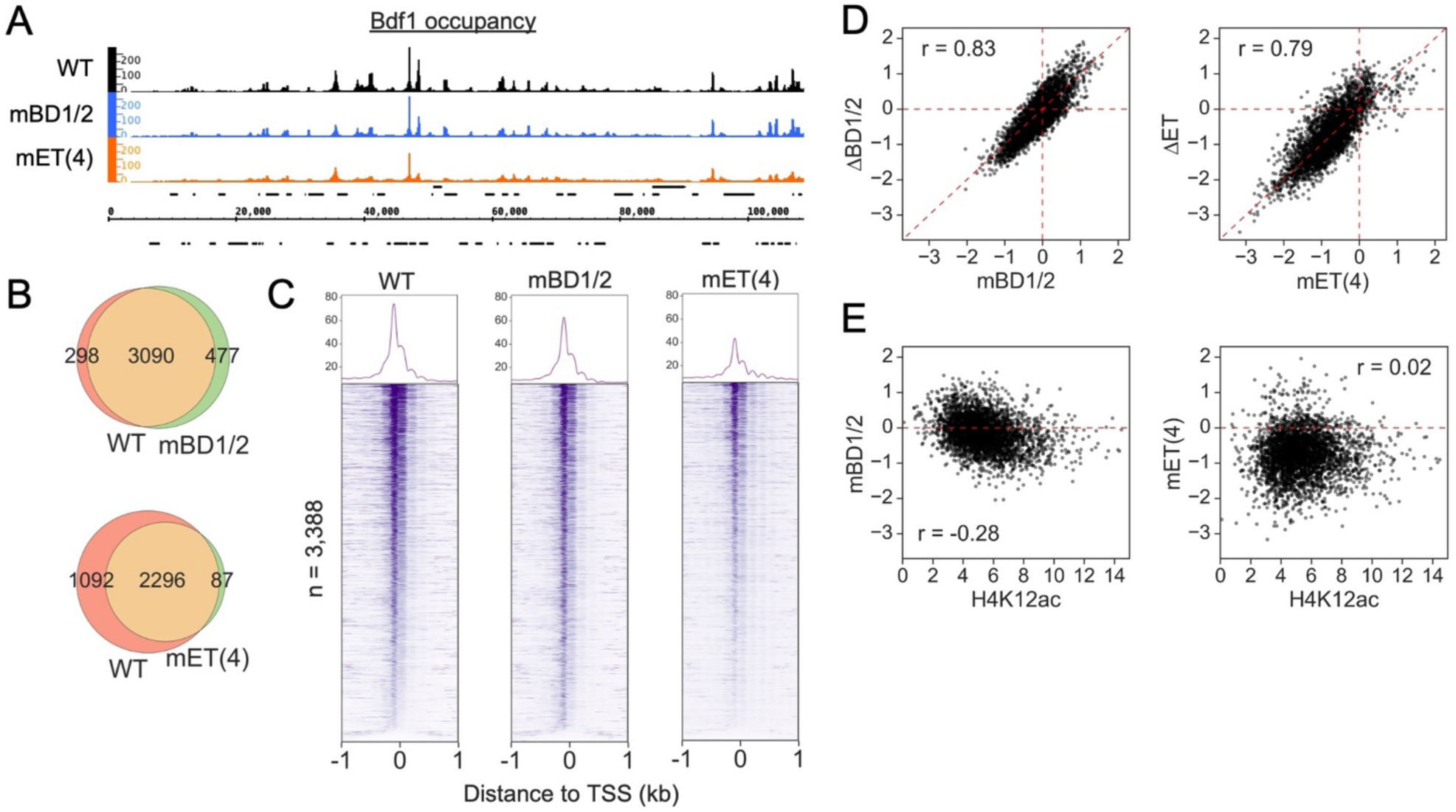
The ET domain has a major role in facilitating BET protein recruitment to chromatin. (**A**) Genome browser snapshot showing the binding patterns of unmodified Bdf1 (WT) and Bdf1 variants with mutations in the BDs or the ET domain as determined by ChEC-seq. Data for the first 100,000 bases of chromosome III are shown. (**B**) Overlap of Bdf1-bound promoters for the indicated experiments. (**C**) Occupancy of unmodified Bdf1 and Bdf1 variants with mutations in the BDs or the ET domain around the transcription start sites (TSS) of 3,388 genes whose promoters are bound by unmodified Bdf1. Genes are sorted by the TSS signal (+/-200 bp) in the WT experiment. (**D**) Comparison of log2 changes in Bdf1 occupancy due to mutation or deletion of the BDs or the ET domain. Bdf1 occupancy was calculated as the sum of signal within the (-100, 100) bp window around the Bdf1 promoter peak summit. Pearson correlation coefficient (r) is shown (*n* = 3,388). (**E**) Comparison of H4K12ac level at promoters ^2^ and log2 changes in Bdf1 occupancy due to mutations in the BDs or the ET domain. Pearson correlation coefficient (r) is shown.

### The ET domain supports TFIID recruitment to gene promoters

Both yeast and mammalian BET proteins have been linked to the coactivator TFIID, which regulates transcription initiation^2,14,15^. Our prior work demonstrated that Bdf1/2 are required for transcription initiation, and that loss of Bdf1/2 modestly decreases TFIID promoter occupancy^2^. Building on this background and our new findings, we hypothesized that a major function of the ET domain in transcription regulation is to facilitate TFIID recruitment. To test this hypothesis, we applied ChEC-seq to quantify changes in TFIID occupancy in cells relying solely on Bdf1 variants after rapid depletion of endogenous Bdf1/2. We inserted an MNase coding sequence into the TFIID subunit Taf1 in three strains previously used in the SLAM-seq assays: the control strain carrying an unmodified additional copy of *BDF1*, and strains carrying *BDF1* variants with mutations in the ET domain or the BDs (mET(4) and mBD1/2, respectively). After depleting endogenous Bdf1/2, we quantified changes in Taf1 recruitment by comparing strains expressing *BDF1* variants to the control strain. Experiments were performed in three biological replicates, and the replicate datasets were highly reproducible (**Fig. S7A**). We found that both the ET domain and the BDs contributed to TFIID recruitment to promoters; however, the ET domain contributed more prominently at most loci (**Fig. 4A-B**). Next, we compared changes in TFIID occupancy resulting from Bdf1 mutations with changes in transcription and observed only a modest correlation (**Fig. S7B**). Consistent with this finding, we previously reported that transcription is uncoupled from baseline TFIID promoter occupancy, and that changes in TFIID occupancy due to complete loss of BET proteins do not correlate with transcriptional output^2,35^. In summary, our results demonstrate that the ET domain plays a major role in TFIID recruitment to gene promoters.

**Fig. 4.**
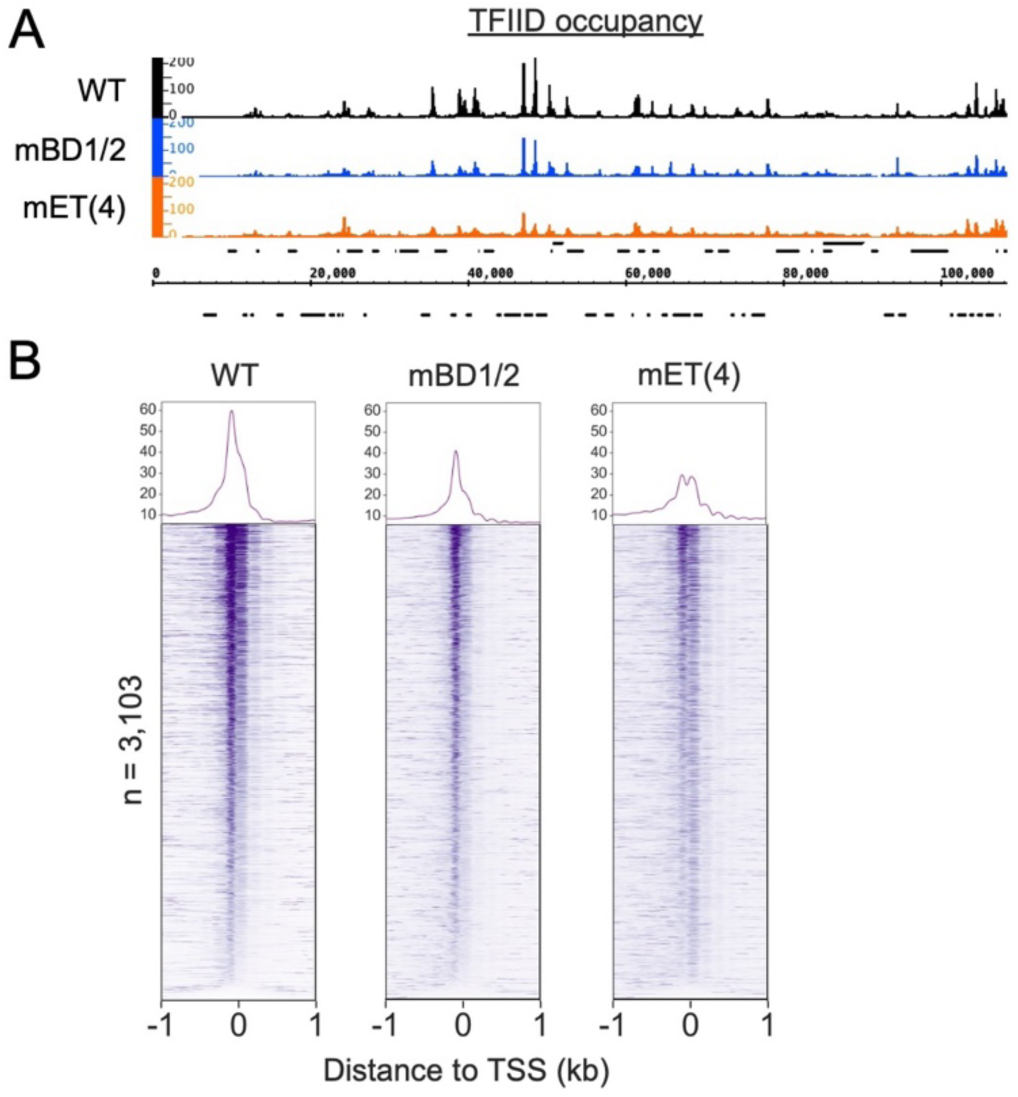
The ET domain facilitates recruitment of TFIID to promoters. (**A**) Genome browser snapshot showing Taf1 (TFIID) binding pattern in the context of unmodified Bdf1 or Bdf1 variants with mutations in the BDs or the ET domain as determined by ChEC-seq. Data for the first 100,000 bases of chromosome III are shown. (**B**) Taf1 occupancy around TSS of 3,103 genes whose promoters are bound by Taf1^2^, in the context of unmodified Bdf1 or Bdf1 variants with mutations in the BDs or the ET domain. Genes are sorted by the signal around TSS (+/-200 bp) in the WT experiment.

### The ET domain engages its targets using a conserved mechanism

Results described thus far show that the ET domain is a primary functional module of Bdf1 and that its impact on transcription is at least partially explained by its role in facilitating TFIID recruitment. Consistent with the latter finding, the ET domain of BRD4 has been proposed to modulate transcription by mediating interactions with protein partners^17,22,23^. However, only a limited number of interactions involving the ET domain of mammalian BET proteins have been experimentally validated, and it remains unknown how many factors associate with the ET domain or what the functional consequences of those interactions are. To begin addressing these questions, we used chromatin mass spectrometry (chromatin-MS) to identify candidate factors – beyond TFIID – that may depend on Bdf1/2 for recruitment to chromatin. We obtained reliable data for 724 of 2399 yeast nuclear proteins and found that 78 exhibited decreased chromatin association following rapid Bdf1/2 depletion. As validation of our approach, this list included two TFIID subunits. Notably, we also identified additional proteins implicated in transcription regulation, including Bur1/CDK9, Spt6, the PAF1 complex (PAF1C) subunit Rtf1, subunits of the RSC, INO80, and ISW2 chromatin remodeling complexes, and several factors involved in mRNA processing (**Figs. 5A and S8A and Table S5**).

**Fig. 5.**
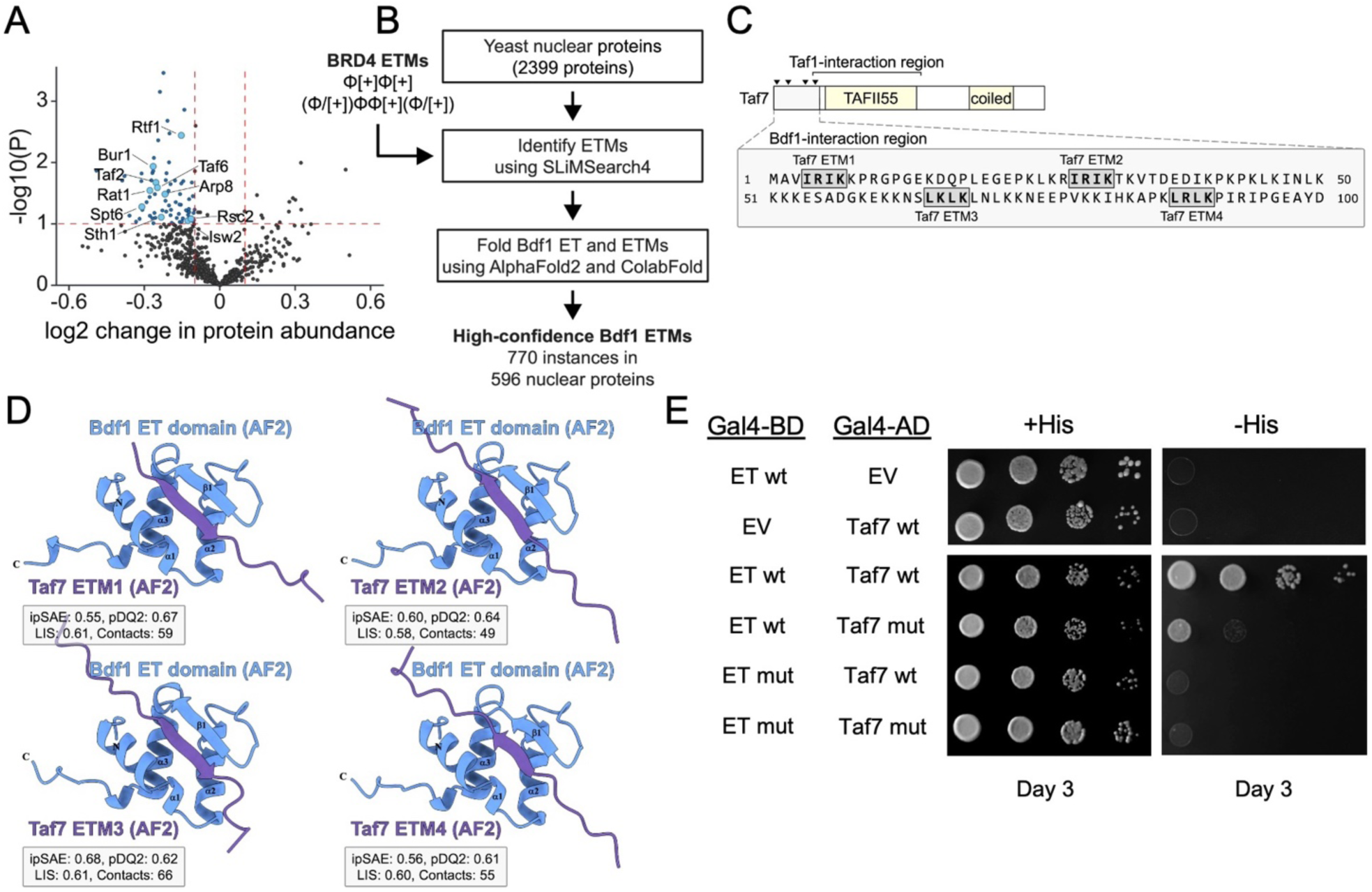
The ET domain of yeast and mammalian BET proteins uses the same mechanism to engage its targets. (**A**) Volcano plot comparing –log10 *p*-value and log2 change in abundance for chromatin-associated proteins following Bdf1/2 depletion. Data for reliably detected nuclear proteins are shown (*n* = 724). (**B**) Schematic representation of the experimental strategy for identifying high-confidence ET-interacting motifs (ETMs) in *S. cerevisiae* nuclear proteins. (**C**) Functional domains of Taf7. Interaction with Bdf1 was previously mapped to the unstructured N-terminal region of Taf7 ^15^, which contains four high-confidence ETMs (detail). The TAFII55 region mediates interaction with Taf1 and is conserved across Taf7 homologs in other eukaryotes. The predicted C-terminal coiled-coil domain is poorly characterized. Other regions of Taf7 are largely unstructured. (**D**) AlphaFold2 (AF2) predicted structures for the Bdf1 ET domain in association with four putative ETMs in the Bdf1-interaction region of Taf7. Confidence metrics (ipSAE, pDockQ2, and LIS) and the number of interfacial contacts supporting structural predictions are shown. (**E**) Y2H analysis of the interaction between the Bdf1 ET domain and Taf7. Plasmids expressing the Bdf1 ET domain (WT or mut), Taf7 (WT or mut), or empty vector (EV) were co-transformed into the host strain. Cells were serially diluted, spotted on indicated media, and imaged after three days of growth (*n* = 3). ET mut – ET domain with four conserved acidic residues substituted with alanine. Taf7 mut – Taf7 with basic residues in the four ETMs in the N-terminal region (first 101 amino acids^15^) substituted with alanine.

Based on proteomic and structural biology studies, the BRD4 ET domain has been proposed to recognize two consensus motifs – Φ[+]Φ[+] or (Φ/[+])ΦΦ[+](Φ/[+]), where Φ is M, L, V, I, or F and [+] is K or R – referred to here as ET-interacting motifs (ETMs)^17,24,25^. Screening the yeast nuclear proteome revealed 2418 ETMs in 1253 proteins, including 60 of the 78 proteins identified in our chromatin-MS experiment (**Fig. 5B and Table S6**). Given this large number of putative interactors, we used AlphaFold2 to model interactions between the Bdf1 ET domain and candidate ETMs^37^. As an initial validation of this approach, we modeled the BRD4 ET domain with selected partner peptides and observed strong similarity between the predicted structures and published experimental structures, including formation of an interfacial β-sheet involving the unstructured loop between the α2 and α3 helices of the ET domain, which adopts a β-strand configuration when bound to partner peptides (**Fig. S8B-C**). Modeling the Bdf1 ET domain with the same peptides similarly produced structures consistent with experimental data, including conditional interfacial β-sheet formation, suggesting a conserved mode of interaction (**Fig. S8D**). We next folded the Bdf1 ET domain with each of the 2418 ETMs identified in yeast nuclear proteins using two implementations of AlphaFold2 and scored models using multiple confidence metrics and interfacial β-sheet establishment (see Materials and Methods) (**Table S7**). To test whether our approach could identify bona fide ET domain interactors, we focused on the TFIID subunit Taf7, a confirmed Bdf1-interacting partner. The interaction was previously mapped to the N-terminal 100 amino acids of Taf7^15^. Within this region, we identified four putative ETMs, each ranking highly in accessibility and model confidence (**Figs. 5C and S8E-F and Table S7**). Structural models of these four ETMs in complex with the Bdf1 ET domain were well supported and showed conditional interfacial β-sheet formation (**Figs. 5D and S9A**). Using a yeast two-hybrid system, we tested whether the Bdf1-Taf7 interaction is mediated by the ET domain and by ETMs in the Taf7 N-terminal region. We found that the Bdf1 ET domain interacts with Taf7 and that this interaction was abolished either by mutating the ET domain or by substituting basic residues with alanine in all four Taf7 ETMs (**Fig. 5E**). Based on these results, together with the observation that conditional β-sheet formation correlates with higher model confidence (**Fig. S9B-C**), we defined 770 high-confidence ETMs (AlphaFold RSA ≥ 0.24 and FoldScore = 1) present in 596 nuclear proteins (**Table S7**). This high-confidence ETM-containing set was enriched for proteins previously associated with Bdf1 function (**Fig. S9D**). Taken together, these findings provide a mechanistic basis for Bdf1-dependent recruitment of TFIID, demonstrate that the ET domains of yeast and mammalian BET proteins engage their targets using a conserved interaction mechanism, and validate our approach for systematically identifying and characterizing interacting partners of the BET protein ET domain.

### Multiple interactions mediated by the ET domain contribute to gene transcription

To further define the mechanistic basis by which the ET domain regulates transcription, we selected the TFIID subunit Taf7 and several additional regulatory factors for detailed analysis. Specifically, we focused on Bur1/CDK9, Spt6, the PAF1C subunit Rtf1, two conserved mRNA-processing factors (Rat1/XRN2 and Rna14/CSTF3), and Pob3, a subunit of the histone chaperone FACT. Each of these proteins contains at least one high-confidence ETM (**Fig. S10 and Table S7**), and all except Pob3 showed decreased chromatin association following Bdf1/2 depletion in our chromatin-MS experiment (**Fig. 5A**). Notably, both FACT subunits Pob3 and Spt16 were previously identified as putative Bdf1-associated proteins by mass spectrometry^18^, and CDK9, FACT, SPT6, PAF1C, XRN2 and CSTF3 have been linked to BET proteins in mammalian cells^5,13,22,38^. However, the mechanistic basis of an association involving BET proteins and any of these factors except CDK9 has not been established.

We first used parallel reaction monitoring mass spectrometry (PRM-MS), a highly quantitative MS method that targets predefined peptides^39^, to measure changes in chromatin association of Bur1, FACT, Spt6, PAF1C, Rat1, and Rna14 following Bdf1/2 depletion. Because Rtf1 could not be reliably detected by PRM-MS, we instead quantified changes in Paf1, the major subunit of PAF1C. For all factors tested, we observed an approximately 50% loss of chromatin association, without a corresponding decrease in total protein levels (**Fig. 6A and Table S5**). Second, we used a yeast two-hybrid system to test interactions between predicted high-confidence ETMs in Bur1, Pob3, and Spt6 (**Fig. S10 and Table S7**) and the ET domain of Bdf1. We found that the Bdf1 ET domain interacted with all predicted targets and that, with the exception of one Pob3 ETM, these interactions were abolished by mutation of either the ET domain or the ETMs (**Fig. S11A**). Based on these results, we generated yeast strains stably expressing variants of Taf7, Bur1, Spt6, or Pob3 in which all identified ETMs were mutated by substituting lysine or arginine residues with alanine. All ETM-mutant strains exhibited moderate growth defects, indicating the functional importance of ET-mediated interactions with the analyzed factors (**Fig. 6B**). Next, we applied SLAM-seq to determine whether ETM-mutant strains display alterations in transcriptional programs (**Table S3**). In the case of the Taf7 and Bur1 mutant strains, we observed downregulation of large subsets of genes (**Fig. 6C**). Conversely, we detected upregulation of many genes in the the Pob3 and Spt6 mutant strains (**Fig. 6C**), suggesting that the interactions between BET proteins and FACT or Spt6 are important for reducing and/or fine-tuning the transcription of certain genes. Consistent with this interpretation, prior studies reported increased coding transcription of subsets of genes upon impaired FACT or Spt6 function^40–42^. Importantly, across all ETM-mutant strains, the subsets of genes that passed the significance threshold overlapped with genes that are strongly sensitive to Bdf1/2 loss (**Fig. S11B**). In summary, our findings demonstrate that ET domain–mediated interactions of BET proteins support chromatin occupancy of multiple partner proteins and enable optimal transcription of target genes.

**Fig. 6.**
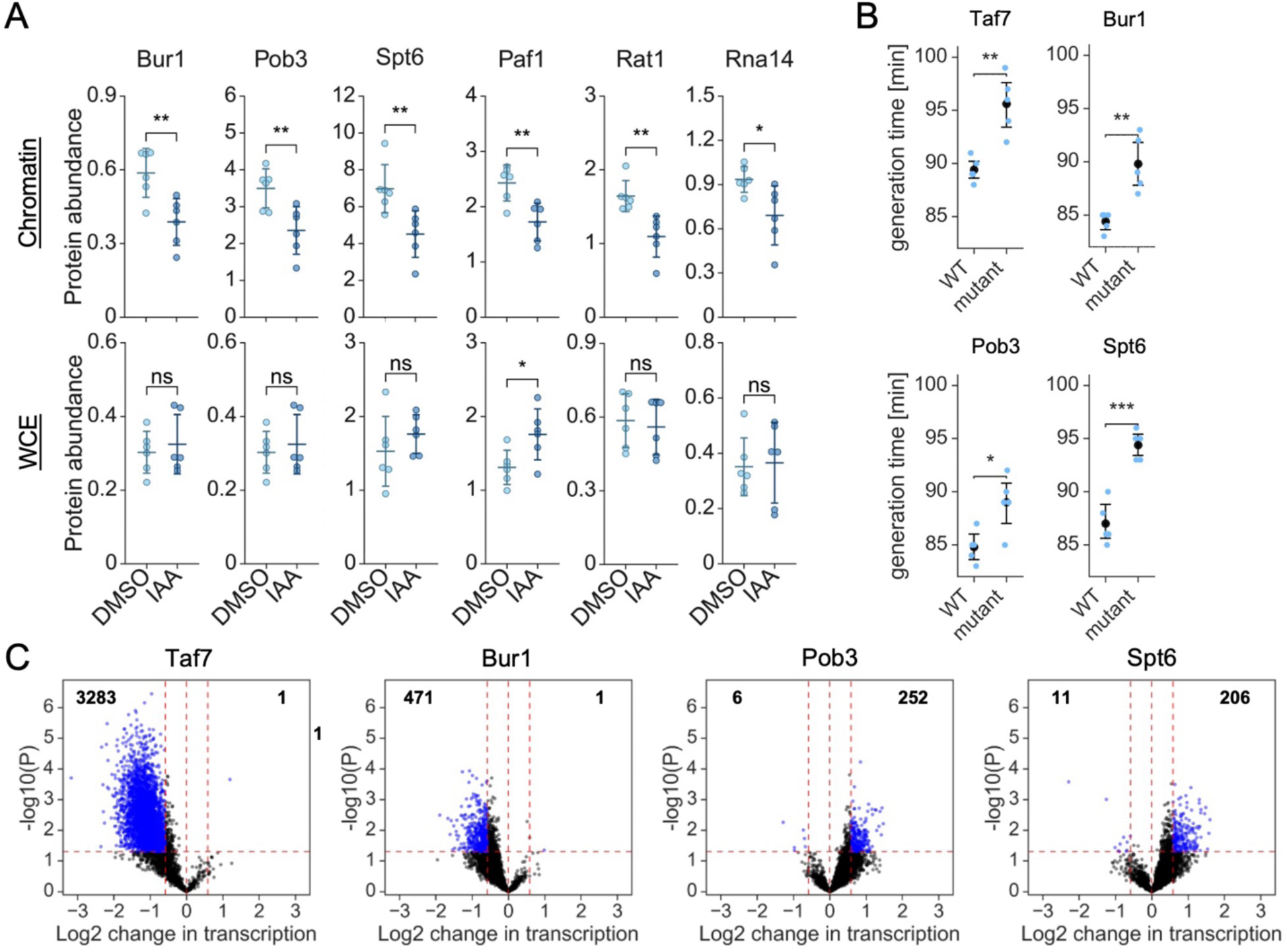
The ET domain modulates transcription by recruiting key regulatory factors. (**A**) Abundance of selected proteins in chromatin-enriched fractions (top panel) or whole cell extract (bottom panel) following Bdf1/2 depletion as determined by parallel reaction monitoring mass spectrometry. Central lines indicate mean protein abundance. Error bars represent standard deviation of the mean (*n* = 6). Results of a Welch’s t-test are shown (* – *p*-value < 0.05, ** – *p*-value < 0.01, ns – not significant). (**B**) Growth assays comparing the fitness of strains expressing variants of indicated proteins carrying targeted ETM mutations to control strains expressing a wild-type variant of the protein from the same genomic locus. Black markers indicate mean generation times. Error bars represent 95% confidence interval (*n* = 5). Results of a Welch’s t-test are shown (* – *p*-value < 0.05, ** – *p*-value < 0.01, *** – *p*-value < 0.001). (**C**) Volcano plots showing log2 change in transcription due to mutation of ETMs in indicated proteins (*n* = 4,498). The numbers of genes showing significant up- or down-regulation are indicated. Thresholds used: fold change – 1.5-fold; *p*-value – 0.05 (Welch’s t-test).

## Discussion

Discovery of specific inhibitors of BET BDs brought the promise of effective treatments for cancer and other diseases. However, the clinical application of BD inhibitors has faced significant challenges due to emerging resistance, cellular heterogeneity, and an incomplete understanding of BET protein biology. In this work, we used a highly versatile yeast model system to define the contributions of all conserved BET protein domains to cell viability, gene transcription, and BET chromatin occupancy. We found that the ET domain, rather than the BDs, constitutes the primary functional module of BET proteins. These results both complement and challenge existing models of BET-mediated transcription, reconcile conflicting observations from prior studies, and suggest new directions for future research.

Many studies have reported BET protein recruitment and function that persist despite BD inhibition^5,10–12,22^. Through genetic analysis, we identified five BET protein domains essential for cell viability, including the BDs, the ET domain, and the phosphorylated NPS/CPS regions. Although the requirement for at least one BD and for the NPS/CPS regions was previously established^18,32^, our work is the first to demonstrate that the ET domain is essential for cell survival. Consistent with the viability defects, we found that the ET domain makes major contributions to global transcription and to BET protein recruitment to chromatin, exceeding the contributions of the BDs. In addition, guided by prior studies^23–25^, we identified conserved acidic residues required for ET domain function *in vivo*. The ET domain has been proposed as a feasible drug target for modulating aberrant transcription in human disease^17,28,29^. Structural information for the ET domain in complex with selected peptide ligands is available, and our data show that the mechanistic basis of ET-mediated interactions is conserved between yeast and mammals. Thus, the yeast model system can serve as a valuable platform to evaluate the ability of small-molecule inhibitors to disrupt ET domain function. Importantly, relative sensitivity of cancer cells to BD inhibition or BET protein degradation differs depending on the tissue of origin and other characteristics, suggesting a possibility of tailored therapeutic approaches^4,8,10^. Altogether, our findings provide a framework and rationale for developing approaches to modulate BET protein activity by targeting the ET domain either in place of, or in combination with, BD inhibition.

Other phenotypes we observed for Bdf1 mutants also complement and extend prior studies. Motif B and the BID were shown to facilitate BRD4 dimerization in biochemical assays, and deletion of motif B caused dissociation of BRD2 from chromatin^31,43^. We observed a significant growth phenotype correlated with changes in Bdf1 recruitment and, to a lesser extent, changes in transcription resulting from deletion of the B/BID regions. Together with prior studies, these findings indicate that BET proteins function as dimers, and future work should address how dimerization modulates BET protein recruitment and activity. For example, an important regulatory layer may arise from the relative ability of BET proteins to form homo- and heterodimers. Phosphorylation of BET proteins in the NPS and CPS regions has been linked with resistance to BD inhibition, interactions with protein partners, and the propensity of BET proteins to dimerize^10,11,30,31^. It was proposed that NPS/CPS phosphorylation may act as a switch that modulates BET protein dimerization and affinity for distinct partners^30,31^. In contrast, our finding that a phosphomimetic *BDF1* variant supports normal growth and causes only mild changes in transcription and Bdf1 chromatin occupancy suggests that dynamic regulation of BET protein phosphorylation is not required for BET protein function. Moreover, prior work proposed distinct roles for the NPS and CPS regions, with NPS phosphorylation linked to BD activity and CPS phosphorylation suggested to influence ET domain function^10,30^. We found that mutation or deletion of NPS did not impair BET protein function, mutation or deletion of CPS was detrimental, and simultaneous perturbation of both regions was lethal. Based on these observations, we propose that phosphorylation of the NPS and CPS regions supports the same functional role or roles of BET proteins. Nevertheless, the biological significance of BET protein phosphorylation remains unresolved and warrants further investigation.

Two recent studies provided mechanistic explanations for BD-independent functions of mammalian BET proteins in transcription^5,22^. Zheng *et al.*^5^ demonstrated that the C-terminal fragment specific to the long isoform of BRD4 can independently promote the release of paused RNA polymerase II (Pol II) through interaction with CDK9/CCNT1 (the P-TEFb complex) and can rescue transcription of a subset of genes following depletion of endogenous BRD4. Yeast lack an equivalent of the long BRD4 isoform, and transcription in yeast does not involve a regulated Pol II pause-and-release mechanism. Our experiments demonstrated that the Bdf1 ET domain interacts with the CDK9 homolog Bur1, and that mutation of Bur1 ETMs impairs cell growth and transcription of a subset of genes. As additional validation, we previously reported defects in Pol II processivity at many of the same genes following Bdf1/2 depletion^2^. Taken together, these observations support a conserved association between BET proteins and Bur1/CDK9 in yeast and mammals, even though the mechanistic basis of the interaction differs. In yeast, the roles of BET proteins in transcription initiation and elongation are mediated primarily by a single Bdf1 isoform and likely by the same functional domain, the ET domain. In mammalian cells, evolutionary acquisition of the long BRD4 isoform may reflect the need to separate BRD4 functions in transcription elongation from other activities also mediated by the short BRD4 isoform, which may be shared with BRD2 and BRD3^14,17,23^. Why such functional separation became advantageous in the context of more complex gene regulatory networks remains an important question for future investigation.

Zhang *et al.*^22^ reported that BRD4 activates transcription at estrogen receptor binding sites through BD-independent associations with SPT5, SPT6, and PAF1C. Consistent with this finding, we observed that Bdf1/2 depletion reduces chromatin association of Spt6 and PAF1C. Spt5, Spt6, and PAF1C subunits each contain high-confidence ETMs, and mutation of Spt6 ETMs resulted in growth defects and dysregulated transcription at many Bdf1/2-dependent genes. Zhang *et al.*^22^ also showed that association with the transcriptional coactivator Mediator contributes to BD-independent BRD4 recruitment. Importantly, the mechanistic basis of the established association between BET proteins and Mediator remains unresolved. Because several Mediator subunits are present in our high-confidence ETM list, it is plausible that BET proteins directly interact with Mediator via the ET domain. Finally, although Bdf1/2 depletion modestly reduces Mediator chromatin occupancy^2^, Bdf1 chromatin association may also be regulated by interactions with Mediator, consistent with the substantial loss of Bdf1 chromatin occupancy observed upon ET domain mutation.

In summary, we provided evidence that the ET domain has a critical role in transcription by serving as a hub for interactions with multiple protein partners, including the transcription coactivator TFIID, Bur1/CDK9, and the histone chaperones Spt6 and FACT. Our analysis indicates that the repertoire of direct ET domain interactors is likely substantially larger. The experimental framework described here can be applied to identify and functionally characterize ET domain interactions with additional transcription regulators in both yeast and mammalian cells. We propose that multiple protein-protein interactions mediated by the ET domain facilitate recruitment of diverse components of the transcription machinery, as well as recruitment of BET proteins themselves. Importantly, this model is consistent with current views of transcription regulation based on the formation of transcriptional condensates through transient interactions among multivalent proteins, and with the propensity of BET proteins to promote condensate formation at gene regulatory elements^44,45^.

Despite fundamental and clinical relevance, many aspects of the biology of BET proteins remain poorly understood. We revealed critical roles of the BET protein ET domain in sustaining global transcription and facilitating BET protein recruitment to chromatin. Our findings provide a new mechanistic explanation for the failure of BD inhibitors to abolish BET protein chromatin occupancy and function in clinical settings and suggest directions for future investigations toward a comprehensive model of the complex roles of BET proteins in transcription regulation.

## Limitations of the study

First, the experiments described in this study were carried out under optimal growth conditions (rich medium, 30°C). As such, it remains to be determined if the contributions of distinct BET protein domains to transcription vary depending on the nutrient availability, stress, or external signaling cues. Second, although prior evidence suggests that Bdf1 is the primary BET protein in yeast cells^2,15^, we cannot exclude the possibility that Bdf2 plays important and non-redundant roles in transcription that are not addressed by our experiments. For example, Bdf2 may be necessary for gene activation rather than steady-state transcription, analogous to proposed roles for BRD2 and BRD3 in mammalian cells ^46^. Consistent with this idea, our recent study identified the *BDF2* promoter as a target of numerous sequence-specific transcription factors, the *BDF2* gene was shown to be regulated by two distinct RNA degradation mechanisms in response to environmental conditions, and Bdf2 was among a small subset of yeast proteins with an unusually short half-life, suggesting dynamic regulation of *BDF2* expression transcriptionally, post-transcriptionally, and post-translationally^47–49^. Finally, there is a possibility that small amounts of endogenous Bdf1/2 remaining after depletion using the AID system may have contributed to confounding effects in our experiments, potentially biasing our interpretation. However, we consider this unlikely, as we observed highly similar results between growth retardation experiments based on Bdf1/2 depletion and *BDF1/2* gene deletion, in which growth was supported by *BDF1* variants expressed from minichromosomal plasmids.

## Materials and Methods

### Yeast cell growth

All *Saccharomyces cerevisiae* and *Schizosaccharomyces pombe* strains used in this study are listed in Table S2. For strain construction, *S. cerevisiae* strains were grown in YPD medium (1% yeast extract, 2% peptone, 2% glucose, 20 μg/ml adenine sulfate) or in synthetic complete (SC) media (0.17% yeast nitrogen base without ammonium sulfate or amino acids [BD Difco], 0.5% ammonium sulfate, 40 μg/ml adenine sulfate, 0.6 mg/ml amino acid dropout mix, supplemented with 2 μg/ml uracil and 0.01% other amino acids to complement auxotrophic markers) at 30°C. Standard amino acid dropout mix contains 2 g each of Tyr, Ser, Val, Ile, Phe, Asp, and Pro and 4 g each of Arg, Thr, Lys, and Met. 1.5% Bacto-agar (BD Difco) was added to medium as required for growth on solid media. For complementation assays, *S. cerevisiae* strains were grown in SC medium lacking Leu to select for cells carrying a minichromosomal plasmid with *BDF1* variants and plated onto medium containing 5-fluoroorotic acid (5-FOA) to shuffle out the wild-type *BDF1* plasmid carrying the URA3 marker. For genomic and growth assays, *S. cerevisiae* cells were grown in YPD medium at 30°C with shaking. *Schizosaccharomyces pombe* cells were grown in YE medium (0.5% yeast extract, 3% glucose) at 30°C with shaking. In experiments involving degron-mediated depletion of target proteins, cells were treated with 1 mM IAA dissolved in DMSO, or with DMSO alone, for 25 min or 3 h to induce protein degradation, followed by protocol-specific steps. For SLAM-seq and ChEC-seq experiments, *S. cerevisiae* cells were collected between an OD600 of 0.5 and 0.7, and *S. pombe* cells were collected at an OD600 of 1.0. For growth assays, cells were grown from an OD600 of 0.0001–0.0005 for 15 h with or without addition of IAA or DMSO. A minimum of five biological replicates were collected for growth assays using strains expressing *BDF1* variants from a minichromosomal plasmid. Three biological replicates were collected for all other experiments unless otherwise indicated.

### Plasmid and strain construction

Plasmids and *S. cerevisiae* strains were constructed using standard methods and are described in **Tables S1 and S2**, respectively. Previously described plasmids or strains were used as indicated^36,50–52^. Proteins were chromosomally tagged by yeast transformation and homologous recombination of PCR-amplified DNA. For ChEC-seq experiments, proteins were tagged with 3xFLAG-MNase::TRP1 or 3xFLAG-MNase::HYG using pGZ110^36^ or pMD75 (this work) plasmids, respectively.

### ET domain overexpression

The gene fragment encoding the Bdf1 ET domain was cloned into the pMD1 plasmid under the control of the *GPM1* promoter and the *BDF1* terminator and placed in frame with the *RPL25* nuclear localization signal and a 3xV5 epitope. The resulting plasmid pJR15 was integrated into the *TRP1* locus of strain RDY73 (Bdf1/2-AID) to generate strain RDY388.

### Western blot analysis

A 1 ml fraction of culture was collected and pelleted from strains after treatment with IAA or DMSO, washed with 500 μl water, and resuspended in 100 μl of yeast whole cell extract buffer (60 mM Tris-HCl [pH 6.8], 10% glycerol, 2% SDS, 5% 2-mercaptoethanol, 0.0025% bromophenol blue). After heating at 95°C for 5 min, samples were centrifuged at 21,000 × g for 5 min and analyzed by SDS-PAGE and western blot. Protein signals were visualized using the Odyssey DLx scanner (Li-Cor) and quantified using the Odyssey Empiria Studio program (Li-Cor). Fluorescent signal from total protein staining (Li-Cor Total Protein Stain workflow) was used to normalize signals for target proteins.

### SLAM-seq

SLAM-seq was performed as previously described^34^. Briefly, following DMSO or IAA treatment, *S. cerevisiae* cultures were treated with 5 mM 4-thiouracil (4tU) in DMSO or with DMSO alone for 4 min. Cells were immediately fixed in cold methanol on dry ice and stored at –80°C. The OD600 at the time of collection was ∼0.7. *S. pombe* cultures were treated with 4tU and collected using the same procedure. Three biological replicates were collected for all experiments. RNA purification and DNase I treatment were performed using the Quick-RNA Fungal/Bacterial Miniprep kit (Zymo Research) according to the manufacturer’s recommendations under reducing conditions, as described. Five micrograms of total RNA was used for alkylation. A total of 200 ng of alkylated RNA was used to construct 3′ mRNA sequencing libraries. Libraries were sequenced on a NovaSeq 6000 (Illumina) at the Oklahoma Medical Research Foundation (OMRF) Clinical Genomics Center (Oklahoma City, OK, USA).

### SLAM-seq data analysis

Data (paired-end 150 bp reads) were analyzed as previously described^34^. Preprocessing and processing steps were implemented as a Snakemake workflow (https://github.com/DonczewLab/SLAM-Seq_Analysis). Briefly, reads were preprocessed using fastp (version 0.23.2)^53^ and bbduk (BBMap version 39.06) (https://sourceforge.net/projects/bbmap/). Reads were then aligned and processed using SLAM-DUNK (version 0.4.3)^54^. Total read counts were defined as the number of reads remaining after alignment and filtering in SLAM-DUNK. T>C read counts were defined as the number of reads containing ≥2 T>C conversions. Genes with zero T>C read counts in one or more replicate samples among all 25 min depletion experiments were excluded. Normalization and differential expression analyses were performed on T>C read counts using DESeq2 (version 1.38.3)^55^. Normalization factors were calculated using total read counts for 25 min depletion experiments. Because large changes in total read counts were expected after 3 h depletion, spike-in read counts were used for normalization in those experiments. Genes that did not meet DESeq2 criteria for fold-change and/or *p*-value calculation were excluded from downstream analyses.

### ChEC-seq

ChEC-seq was performed as previously described with minor modifications^2,56,57^. Briefly, 50 ml *S. cerevisiae* cultures were pelleted at 2000 × g for 3 min. Cells were resuspended in 1 ml of Buffer A (15 mM Tris-HCl [pH 7.5], 80 mM KCl, 0.1 mM EGTA, 0.2 mM spermine [Millipore Sigma], 0.3 mM spermidine [Millipore Sigma], 1X protease inhibitors [2 μg/ml aprotinin, 1 μg/ml pepstatin A, 1 μg/ml leupeptin, 2 mM PMSF]), transferred to a 1.5 ml microcentrifuge tube, and pelleted at 1500 × g for 30 s. Cells were washed twice with 1 ml of Buffer A and resuspended in 570 μl of Buffer A. Thirty microliters of 2% digitonin (Millipore Sigma) was added to a final concentration of 0.1%, and cells were permeabilized for 5 min at 30°C with shaking (900 rpm). Samples were incubated with 0.2 mM CaCl2 for 5 min at 30°C with shaking. A 100 μl fraction of the suspension was mixed with 100 μl of Stop Solution (400 mM NaCl, 20 mM EDTA, 4 mM EGTA). Samples were then incubated with 0.4 mg/ml Proteinase K (Thermo Fisher Scientific) for 30 min at 55°C. DNA was purified using the ChIP DNA Clean and Concentrator kit (Zymo Research) and eluted in 30 μl of elution buffer. Samples were treated with 0.3 mg/ml RNase A (Thermo Fisher Scientific) for 20 min at 37°C. Thirty microliters of Sera-Mag Speedbeads (Cytiva), prepared as described (dx.doi.org/10.17504/protocols.io.x54v9p7b1g3e/v2), were added, and samples were incubated for 10 min at room temperature. Supernatants were transferred to a new tube, and the volume was adjusted to 175 μl (10 mM Tris-HCl, pH 8.0). DNA was purified again using the ChIP DNA Clean and Concentrator kit and eluted in 30 μl of Elution Buffer. Libraries were prepared using the Next Ultra II DNA Library Prep kit (New England Biolabs) as described^58^ and sequenced on a NovaSeq 6000 at the OMRF Clinical Genomics Center.

### ChEC-seq data analysis

Data (paired-end 150 bp reads) were analyzed as previously described with minor modifications^2,56^. Initial preprocessing and processing steps were implemented as a Snakemake workflow (https://github.com/DonczewLab/ChEC-Seq_Analysis). Briefly, reads were preprocessed using bbduk (BBMap version 39.06) (https://sourceforge.net/projects/bbmap/) and aligned using Bowtie 2 (version 2.5.0)^59^. BAM files were generated, sorted, and indexed using Samtools (version 1.18)^60^. Genome coverage was calculated and normalized by counts per million (CPM) using deepTools2 (version 3.5.4)^61^. Average genome coverage across replicates was calculated using BEDTools (version 2.30.0)^62^ and bedGraphToBigWig^63^. SAM files for *S. cerevisiae* were converted to tag directories using the HOMER^64^ ‘makeTagDirectory’ tool. Peaks were called using the HOMER ‘findPeaks’ tool with optional arguments set to ‘-o auto -C 0 L 2 F 2’, using the free MNase dataset as a control^2^. These settings apply a default false discovery rate (FDR; 0.1%) and require peaks to be enriched twofold over the control and twofold over the local background. Resulting peak files were converted to BED format using the ‘pos2bed.pl’ program. For each peak, the summit was defined as the midpoint between the peak borders. For promoter assignment, the list of annotated ORF sequences (excluding entries classified as ‘dubious’ or ‘pseudogene’) was downloaded from the SGD website (https://www.yeastgenome.org). Data for 5888 genes were merged with TSS positions^65^. If TSS annotation was missing, the TSS was manually assigned at −100 bp relative to the start codon. At this stage, biological replicates were averaged for each sample. Peaks were assigned to promoters if their summit was located between −300 and +100 bp relative to the TSS. When more than one peak mapped to a promoter, the peak closest to the TSS was used. Promoters bound in at least two out of four replicate experiments were included in the final list. To quantify log2 changes in promoter occupancy, signal per promoter was calculated as the sum of normalized reads within a 200 bp window centered on the promoter peak summit.

### ET-interacting motif discovery

Putative ET-interacting motifs (ETMs) were identified using SLiMSearch (version 4) ^66^. The previously described ETMs Φ[+]Φ[+] and (Φ/[+])ΦΦ[+](Φ/[+]), where Φ is M, L, V, I, or F and [+] is K or R^17^, were used as input to query the *S. cerevisiae* proteome. ETMs were limited to those present in nuclear proteins based on evidence in AllianceMine^67^. Overlapping and/or adjacent ETMs were collapsed. Disorder propensity and accessibility were calculated in SLiMSearch using IUPred2A^68^ and AlphaFold RSA^69^, respectively.

### Structural predictions

Initial folding experiments involving characterized BRD4-interacting peptides and the BRD4 ET domain or the Bdf1 ET domain were performed using a locally installed instance of AlphaFold2 (version 2.3.2)^37,70^. Experimental structures were obtained from the Protein Data Bank (PDB)^71^. ChimeraX (version 1.10.1)^72^ was used to visualize structures and calculate structural similarity and interfacial contacts. Similarity between experimental and predicted structures was calculated as root mean square deviation (RMSD). Contacts were calculated as the number of interfacial residue pairs ≤ 4.0 Å apart with a maximum predicted aligned error (PAE) ≤ 5.0 Å. ETMs in nuclear proteins (*n* = 2418) were folded with the Bdf1 ET domain using two implementations of AlphaFold2 : the locally installed AlphaFold2 instance (as above) and ColabFold Batch (version 1.5.5)^73^ run on Google Colaboratory. Eight amino acids N-terminal and eight amino acids C-terminal to each ETM were included in folding experiments. Confidence metrics were calculated using the ipSAE program (version 3)^74^. Cutoffs of PAE ≤ 10 Å and distance ≤ 10 Å were used to compute ipSAE scores. Combined ipSAE confidence was calculated as the unweighted z-score average (scaled 0-1) across ipSAE scores of the top-ranked AlphaFold2 and ColabFold models. Combined pDockQ2 confidence was calculated as the unweighted z-score average (scaled 0-1) across pDockQ2 scores of the top-ranked AlphaFold2 and ColabFold models. The DSSP program (version 4.5.6)^75^ was used to assign secondary structure elements (i.e., interfacial β-sheet establishment) in predicted structures. FoldScore was calculated based on conditional interfacial β-sheet establishment across top-ranked AlphaFold2 and ColabFold models: a FoldScore of 1 indicates that interfacial β-sheets with the same orientation were assigned in both models; a FoldScore of 0.5 indicates that interfacial β-sheets with different orientations were assigned in the two models or that an interfacial β-sheet was assigned in only one model; and a FoldScore of 0 indicates that an interfacial β-sheet was not assigned in either model.

### Chromatin mass spectrometry

Chromatin-enriched fractions and whole cell extract were collected using chromatin-enriched fractionation (ChEF) as previously described, with modifications^76^. Briefly, *S. cerevisiae* was grown in 200 ml YPD medium at 30°C with shaking at 220 rpm to an OD600 of ∼0.5 and split into two 100 ml cultures. Cultures were treated with IAA or DMSO for 25 min. Cells were pelleted at 2000 × g for 3 min, washed once with sterile water, and stored at –80°C. The OD600 at the time of collection was ∼0.7. A minimum of five biological replicates were collected. Cells were resuspended in 1 ml ChEF buffer 1 ^76^ supplemented with 1X phosphatase inhibitors (Apex Biotechnology), 1X deacetylase inhibitors (Apex Biotechnology), and 1X protease inhibitors (see above). Cells were combined with 1.5 ml disruption beads and processed in a Mini-Beadbeater-24 (Biospec Products) at 3800 rpm for 30 s followed by incubation on ice for 2 min, repeated 10 times. Cell lysate was recovered as previously described^77^ and cleared twice by centrifugation at 500 × g for 5 min at 4°C. A fraction of the cleared lysate represented whole cell extract. The remaining lysate was processed as described previously^76^, except that ChEF buffer 2 was supplemented with 1X phosphatase inhibitors, 1X deacetylase inhibitors, and 1X protease inhibitors. Chromatin-enriched fractions were resuspended in 50 mM Tris-HCl (pH 7.5).

A total of 15 pmoles of bovine serum albumin (BSA) was added to 100 μg of total protein from chromatin-enriched fractions or 200 μg of total protein from whole cell extracts as an internal standard. Samples were processed as previously described^78^. Samples from chromatin-enriched fractions were analyzed using a Q Exactive Plus mass spectrometer system (ThermoFisher Scientific) in data-independent acquisition (DIA) mode with a 20 m/z window. Samples from whole cell extracts were analyzed using an Exploris 480 mass spectrometer system (ThermoFisher Scientific) in DIA mode with a 10 m/z window. Data were analyzed using DIA-NN (version 1.9.2)^79^ searching against the *S. cerevisiae* UniProt protein database (downloaded from ENA/EMBL, accession GCA_000146045.2).

### Parallel reaction monitoring mass spectrometry

Whole cell extracts and chromatin-enriched fractions were collected as described above. A total of 14 pmoles of BSA was added to 60 μg of total protein from chromatin-enriched fractions or 100 μg of total protein from whole cell extracts as an internal standard. Samples were processed as previously described^78^ and analyzed using a Stellar mass spectrometer system (ThermoFisher Scientific) in PRM mode with a resolution of 1.2 Da and a scan rate of 67 kDa/sec. Data were processed using Skyline (version 25.1)^80^ to identify and integrate chromatographic peaks for a defined set of validated peptides. Retention times were calibrated using BSA and trypsin peptides. Protein abundance was calculated as the geometric mean of a minimum of two monitored peptides normalized to the BSA internal standard and expressed as pmol per 10 µg of total protein.

### Yeast two-hybrid analysis

Yeast two-hybrid experiments were performed using the Matchmaker Gold Y2H system (Takara Bio) based on the manufacturer’s recommendations with modifications. Briefly, the *S. cerevisiae* strain Y2HGold was co-transformed with *TRP1*-marked Gal4-BD and *LEU2*-marked Gal4-AD plasmids, pGBKT7 and pGADT7, respectively. Co-transformed strains were cultured in SC-Leu-Trp medium at 30°C overnight, adjusted to an OD600 of 1, and serially diluted tenfold. Five microliters of each dilution was spotted onto SC-Leu-Trp, SC-Leu-Trp-His, or SC-Leu-Trp-His plates supplemented with 2 mM 3-Amino-1,2,4-triazole (3-AT, MilliporeSigma). Plates were incubated at 30°C and imaged daily for a minimum of four days.

## Acknowledgments

We thank Linda Thompson, Dean Dawson, Christopher Sansam and Karen Arndt for helpful comments, Steven Hahn for plasmids, and Stephen Buratowski for anti-Bdf1 antibody. We thank the OMRF Research Computing Services, Clinical Genomics Center, and Center for Biomedical Data Sciences for contributions to this work.

## Author contributions

Conceptualization: JR, MD, RD

Methodology: JR, MD, MK, KB, RD

Investigation: IP, JR, AM, MD, MK, RD

Visualization: IP, JR, AM, RD

Supervision: JR, MD, RD

Writing—original draft: RD

Writing—review & editing: JR, MD, RD

## Data and materials availability

All data are available in the main text or the supplementary materials. SLAM-seq and ChEC-seq datasets have been deposited in the NCBI Gene Expression Omnibus (GEO) under accession number GSE313013. Raw datasets from DIA and PRM mass spectrometry experiments have been deposited in the Zenodo repository (https://doi.org/10.5281/zenodo.17966389). Reference genome assemblies and annotations for *S. cerevisiae* strain S288C (version R64-3-1) and *S. pombe* strain 972h- (version ASM294v2) were retrieved from the NCBI Datasets repository (GCF 000146045.2 and GCF 000002945.1, respectively). BED files for *S. cerevisiae* and *S. pombe* used in SLAM-DUNK were retrieved from the Zenodo repository (https://doi.org/10.5281/zenodo.10714018).

