## Supplementary Information for "The extra-terminal domain drives the role of BET proteins in transcription"

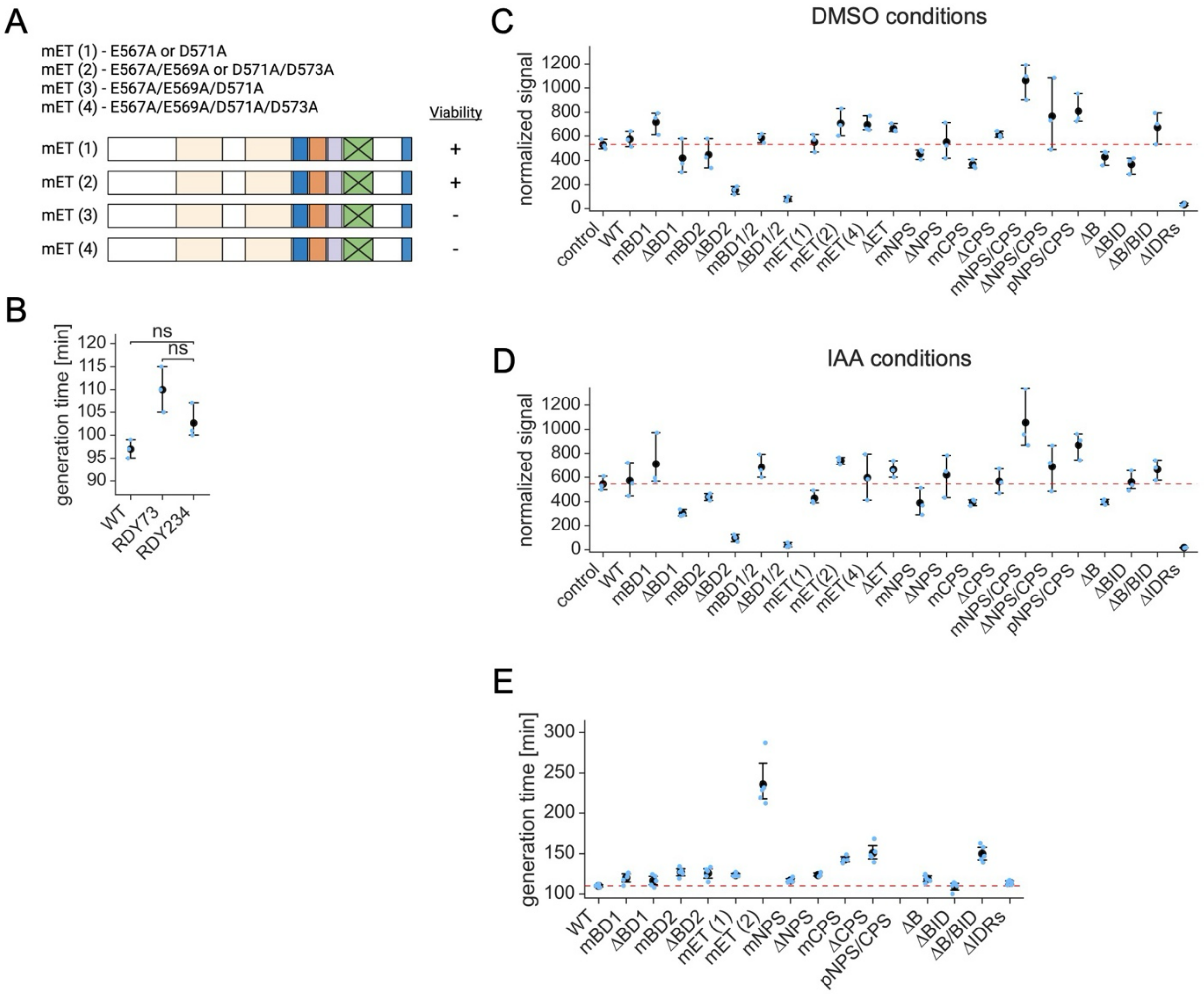
A
mET (1) - E567A or D571A
mET (2) - E567A/E569A or D571A/D573A
mET (3) - E567A/E569A/D571A
mET (4) - E567A/E569A/D571A/D573A
Viability
mET (1)
+
mET (2)
+
mET (3)
-
mET (4)
-
B
generation time [min]
ns
ns
WT
RDY73
RDY234
C
DMSO conditions
normalized signal
control
WT
mBD1
ΔBD1
mBD2
ΔBD2
mBD1/2
ΔBD1/2
mET(1)
mET(2)
mET(4)
ΔET
mNPS
ΔNPS
mCPS
ΔCPS
mNPS/CPS
ΔNPS/CPS
pNPS/CPS
ΔB
ΔBID
ΔB/BID
ΔIDRs
D
IAA conditions
normalized signal
E
generation time [min]
WT
mBD1
ΔBD1
mBD2
ΔBD2
mET (1)
mET (2)
mNPS
ΔNPS
mCPS
ΔCPS
pNPS/CPS
ΔB
ΔBID
ΔB/BID
ΔIDRs

**Fig. S1. At least two conserved acidic residues in the ET domain are required for viability.** (**A**) Six Bdf1 variants carrying targeted mutations in the ET domain. Ability to support growth in the absence of endogenous *BDF1/2* is indicated. (**B**) Growth assay comparing the fitness of a laboratory wild-type strain (BY4705, WT), the RDY73 strain, and an RDY73 derivative expressing additional copy of unmodified *BDF1* from the *TRP1* locus (RDY234). Black markers indicate mean generation times. Error bars represent the 95% confidence interval ($n$ = 3). Results of a Welch's t-test are shown (ns – not significant). (**C**) Western blot analysis of the expression of the indicated *BDF1* variants from the *TRP1* locus of strain RDY73 without depletion of endogenous Bdf1/2 (DMSO conditions). The reference expression level (red dashed line) corresponds to the parental strain BY4705. An anti-Bdf1 antibody was used. Fluorescent signal for total protein content (LI-COR total protein stain workflow) was used to normalize target proteins. Black markers indicate mean normalized signal. Error bars represent the 95% confidence interval ($n$ = 3). (**D**) Same as (**C)** but with depletion of endogenous Bdf1/2 (IAA conditions) ($n$ = 3). (**E**) Growth assay comparing the fitness of a control strain expressing unmodified *BDF1* (WT) from a minichromosomal plasmid to strains expressing *BDF1* variants in the context of deletion of endogenous *BDF1/2*. Black markers indicate mean generation time. Error bars represent the 95% confidence intervals ($n$ = 4-5).

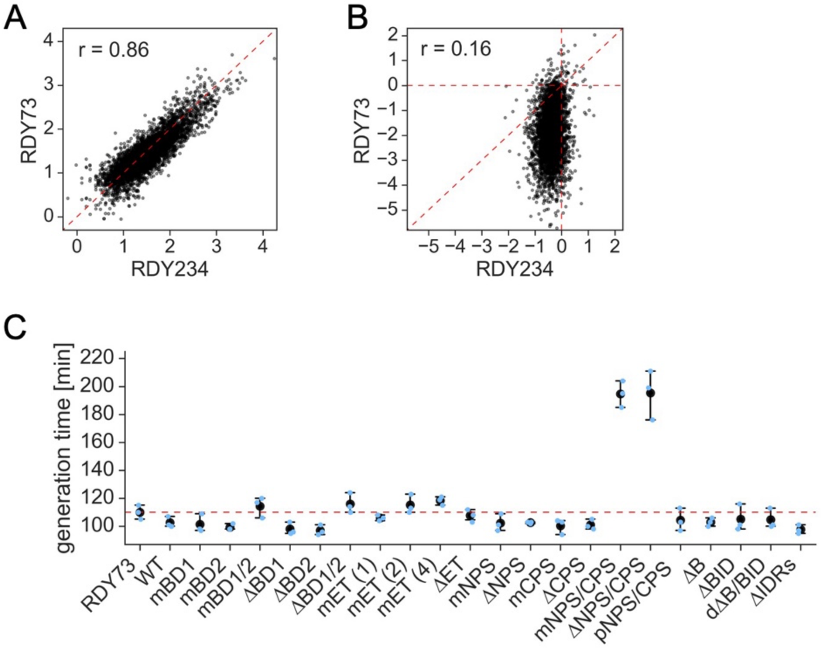
A
r = 0.86
RDY73
RDY234
B
r = 0.16
RDY73
RDY234
C
generation time [min]
RDY73
WT
mBD1
mBD2
mBD1/2
ΔBD1
ΔBD2
ΔBD1/2
mET (1)
mET (2)
mET (4)
ΔET
mNPS
ΔNPS
mCPS
ΔCPS
mNPS/CPS
ΔNPS/CPS
pNPS/CPS
ΔB
ΔBID
dΔB/BID
ΔIDRs

**Fig. S2. Additional copy of *BDF1* is well tolerated by yeast cells.** (**A**) Comparison of baseline transcription ($\log_{10}$) in strains RDY73 (Bdf1/2-AID) and its derivative expressing an additional copy of unmodified *BDF1* from the *TRP1* locus (RDY234). Pearson correlation coefficient (r) is shown ($n$ = 4,836). (**B**) Log2 changes in transcription measured by SLAM-seq after depleting endogenous Bdf1/2 for 25 min in strains RDY73 and RDY234. Pearson correlation coefficient (r) is shown ($n$ = 4,836). (**C**) Growth assay comparing the fitness of strains expressing unmodified Bdf1 (WT) or the indicated Bdf1 variants to the parental strain RDY73 in the presence of endogenous Bdf1/2 (no IAA treatment). Black markers indicate mean generation time. Error bars correspond to 95% confidence interval ($n$ = 3).

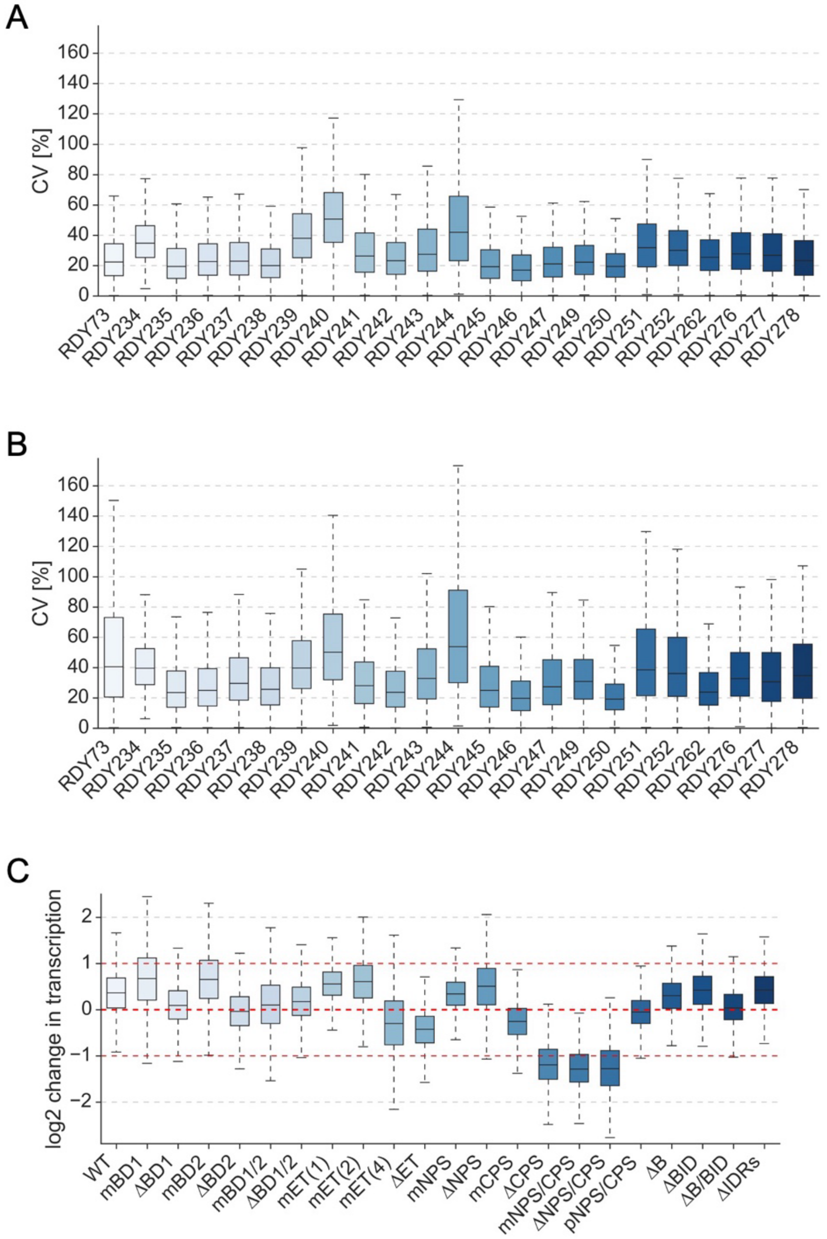
A
CV [%]
0
20
40
60
80
100
120
140
160
RDY73
RDY234
RDY235
RDY236
RDY237
RDY238
RDY239
RDY240
RDY241
RDY242
RDY243
RDY244
RDY245
RDY246
RDY247
RDY249
RDY250
RDY251
RDY252
RDY262
RDY276
RDY277
RDY278
B
CV [%]
0
20
40
60
80
100
120
140
160
RDY73
RDY234
RDY235
RDY236
RDY237
RDY238
RDY239
RDY240
RDY241
RDY242
RDY243
RDY244
RDY245
RDY246
RDY247
RDY249
RDY250
RDY251
RDY252
RDY262
RDY276
RDY277
RDY278
C
log2 change in transcription
2
1
0
−1
−2
WT
mBD1
ΔBD1
mBD2
ΔBD2
mBD1/2
ΔBD1/2
mET(1)
mET(2)
mET(4)
ΔET
mNPS
ΔNPS
mCPS
ΔCPS
mNPS/CPS
ΔNPS/CPS
pNPS/CPS
ΔB
ΔBID
ΔB/BID
ΔIDRs

**Fig. S3. SLAM-seq accurately captures rapid changes in transcription resulting from targeted mutations in BET protein domains.** (**A**) Coefficient of variation (CV) for the indicated SLAM-seq experiments under DMSO conditions. CV values were calculated for each gene based on normalized read counts from replicate experiments (**Table S3**) ($n$ = 4,836). (**B**) Coefficient of variation (CV) for the indicated SLAM-seq experiments under IAA conditions. CV values were calculated for each gene based on normalized read counts from replicate experiments (**Table S3**) ($n$ = 4,836). **(C)** Log2 changes in transcription due to *BDF1* mutations without depletion of endogenous Bdf1/2 ($n$ = 4,836).

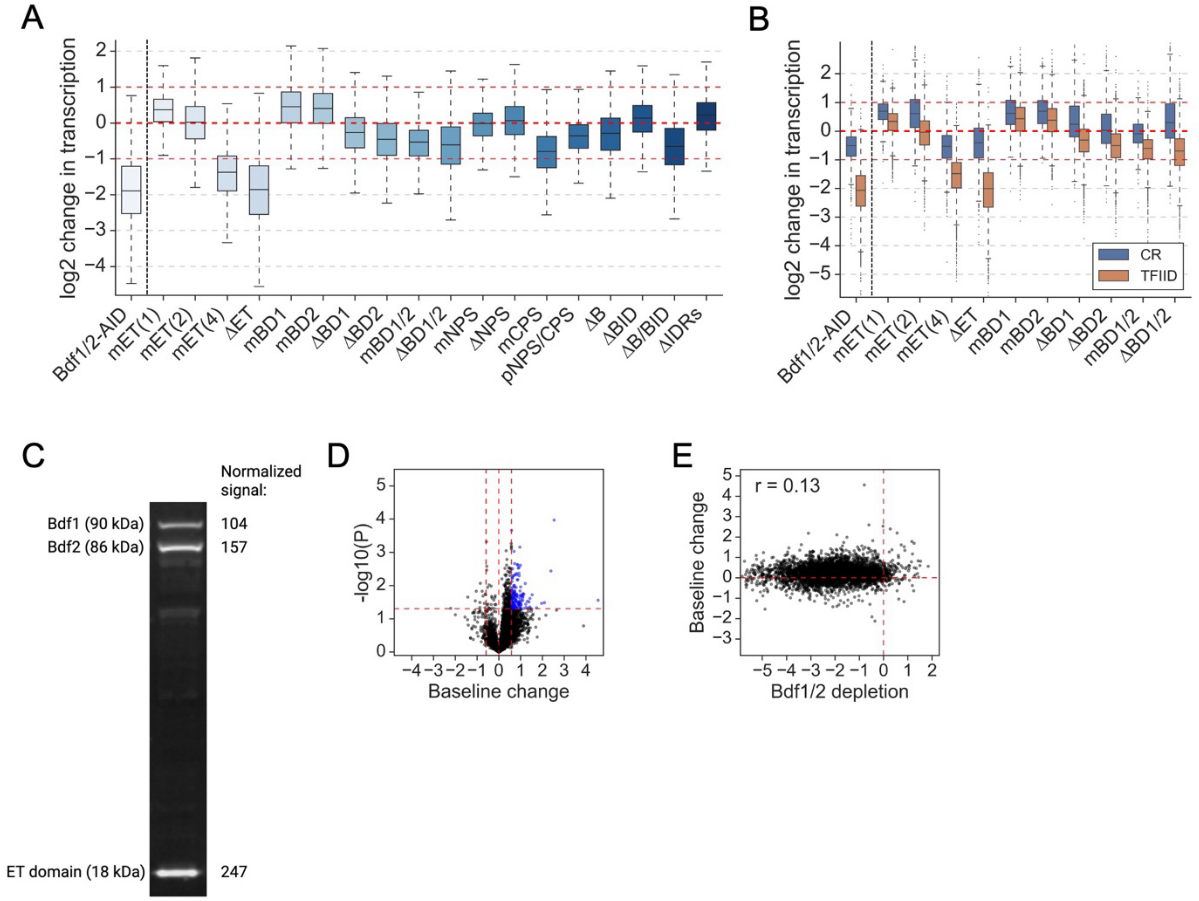
A
B
C
D
E
log2 change in transcription
Bdf1/2-AID
mET(1)
mET(2)
mET(4)
ΔET
mBD1
mBD2
ΔBD1
ΔBD2
mBD1/2
ΔBD1/2
mNPS
ΔNPS
mCPS
pNPS/CPS
ΔB
ΔBID
ΔB/BID
ΔIDRs
CR
TFIID
Normalized signal:
Bdf1 (90 kDa)
Bdf2 (86 kDa)
ET domain (18 kDa)
104
157
247
-log10(P)
Baseline change
r = 0.13
Bdf1/2 depletion

**Fig. S4. The ET domain is the only BET protein domain broadly required for transcription.** (**A**) Log2 changes in transcription due to *BDF1* mutations measured by SLAM-seq after depleting endogenous Bdf1/2 for 25 min (*n* = 4,836). Results of depletion of Bdf1/2 in the absence of an additional copy of *BDF1* are shown in the first column from the left. (**B**) Log2 changes in transcription due to *BDF1* mutations measured by SLAM-seq after depleting endogenous Bdf1/2 for 25 min. Genes are divided into previously defined major yeast gene classes (CR – coactivator-redundant, TFIID – TFIID-dependent) (*35*) (*n* = 4,560). (**C**) Western blot analysis of the ET domain overexpression in the RDY73 derivative strain RDY388. An anti-V5 antibody was used to detect target proteins. Fluorescent signal for total protein content (LI-COR total protein stain workflow) was used to normalize target proteins. A ~13-fold ET domain overexpression compared to Bdf1 was observed after correcting for the size difference. A representative image of three biological replicates is shown. (**D**) Volcano plot comparing transcriptional changes due to ET domain overexpression (log2 scale) in the presence of endogenous Bdf1/2 with the associated *p*-value. Thresholds used: fold change – 1.5-fold, *p*-value – 0.05 (Welch's t-test). (**E**) Comparison of transcriptional changes due to ET domain overexpression in the presence of endogenous Bdf1/2 (y-axis) with gene dependence on Bdf1/2 (x-axis). Pearson correlation coefficient (r) is shown.

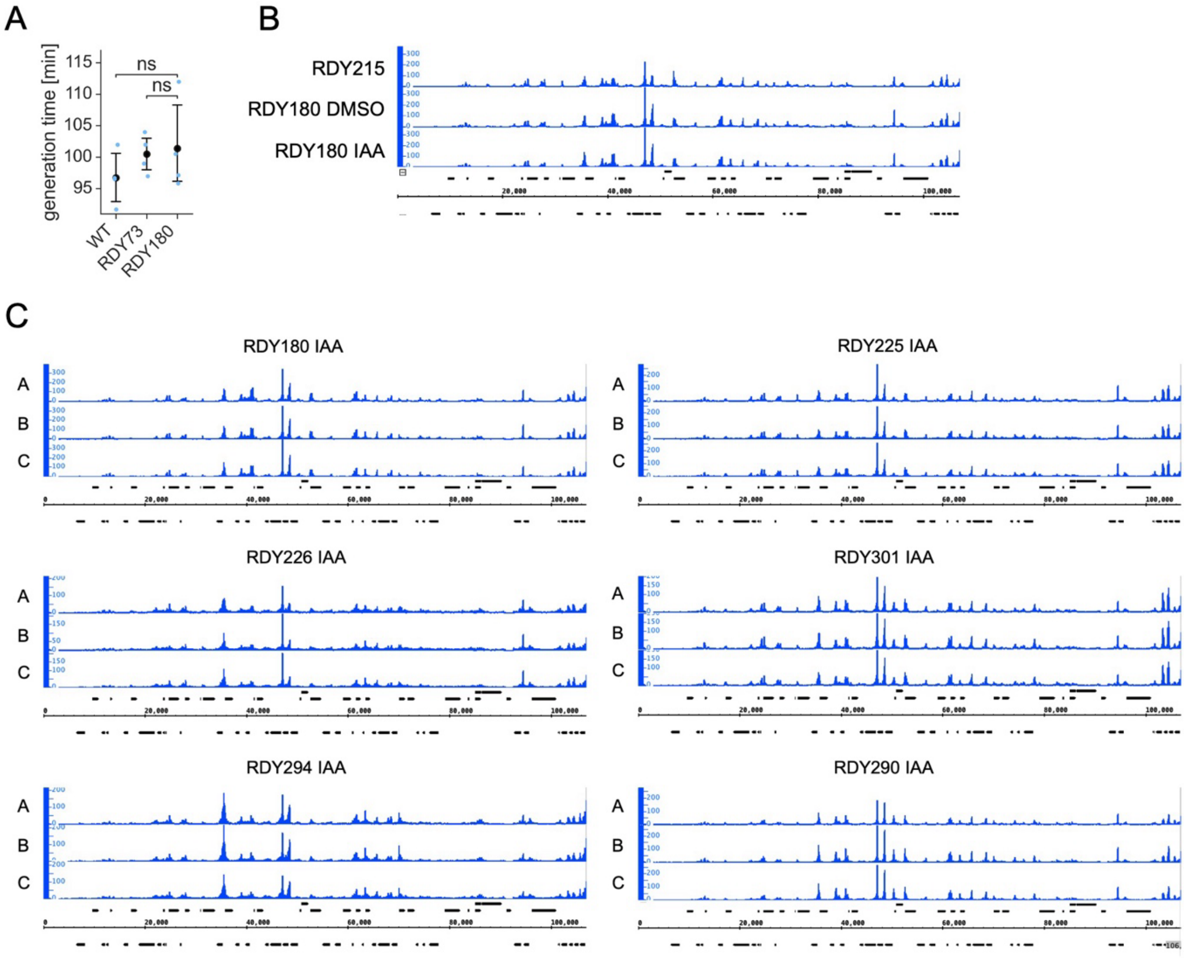
A
B
C
generation time [min]
115
110
105
100
95
ns
ns
WT
RDY73
RDY180
RDY215
RDY180 DMSO
RDY180 IAA
RDY225 IAA
RDY226 IAA
RDY301 IAA
RDY294 IAA
RDY290 IAA
20,000
40,000
60,000
80,000
100,000

**Fig. S5. ChEC-seq allows reproducible mapping of chromatin occupancy for ectopically expressed Bdf1 variants.** (**A**) Growth assay comparing the fitness of a laboratory wild-type strain (BY4705, WT), the RDY73 strain, and a RDY73 derivative strain expressing a *BDF1-MNase* fusion protein from the *TRP1* locus (RDY180). Black markers indicate mean generation time. Error bars correspond to the 95% confidence interval ($n$ = 3). Results of a Welch's t-test are shown (ns – not significant). (**B**) Genome browser snapshot comparing a binding pattern of ectopically expressed Bdf1 in strain RDY180 in the presence (DMSO) or absence (IAA) of endogenous Bdf1/2 to that of the reference strain RDY215. Data for the first 100,000 bases of chromosome III is shown**.** (**C**) Comparison of ChEC-seq data for individual replicate experiments for selected strains. Data for the first 100,000 bases of chromosome III is shown.

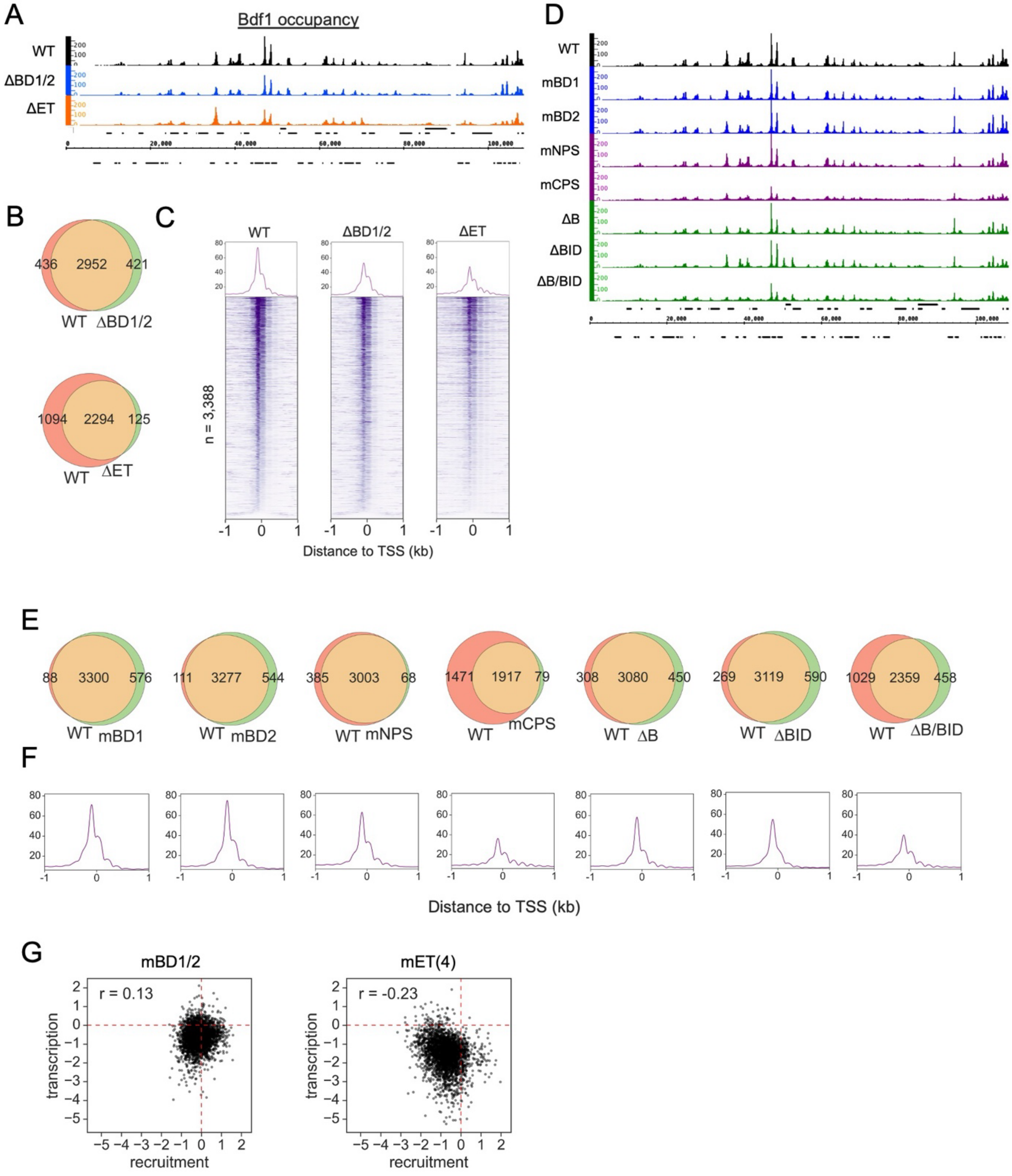
A
Bdf1 occupancy
WT
ΔBD1/2
ΔET
B
436 2952 421
WT ΔBD1/2
1094 2294 125
WT ΔET
C
WT
ΔBD1/2
ΔET
n = 3,388
Distance to TSS (kb)
D
WT
mBD1
mBD2
mNPS
mCPS
ΔB
ΔBID
ΔB/BID
E
88 3300 576
WT mBD1
111 3277 544
WT mBD2
385 3003 68
WT mNPS
1471 1917 79
WT mCPS
308 3080 450
WT ΔB
269 3119 590
WT ΔBID
1029 2359 458
WT ΔB/BID
F
Distance to TSS (kb)
G
mBD1/2
r = 0.13
transcription
recruitment
mET(4)
r = -0.23
transcription
recruitment

**Fig. S6. CPS and B/BID regions are important for Bdf1 chromatin occupancy.** (**A**) Genome browser snapshot showing the binding pattern of unmodified Bdf1 (WT) and Bdf1 variants with deletions of the BDs or the ET domain as determined by ChEC-seq. Data for the first 100,000 bases of chromosome III is shown. (**B**) Overlap of Bdf1-bound promoters for the indicated experiments. (**C**) Occupancy of unmodified Bdf1 and Bdf1 variants with mutations in the BDs or the ET domain around the transcription start sites (TSS) of 3,388 genes whose promoters are bound by unmodified Bdf1. Genes are sorted by the TSS signal (+/- 200 bp) in the WT experiment. (**D**) Genome browser snapshot showing binding pattern of unmodified Bdf1 (WT) and the indicated Bdf1 variants as determined by ChEC-seq. Data for the first 100,000 bases of chromosome III is shown. (**E**) Overlap of Bdf1-bound promoters for the indicated experiments. (**F**) Average occupancy of the indicated Bdf1 variants around the transcription start sites (TSS) of 3,388 genes whose promoters are bound by unmodified Bdf1**.** (**G**) Comparison of changes in Bdf1 promoter occupancy due to mutations in the BDs or the ET domain and corresponding changes in transcription. Pearson correlation coefficient (r) is shown.

A

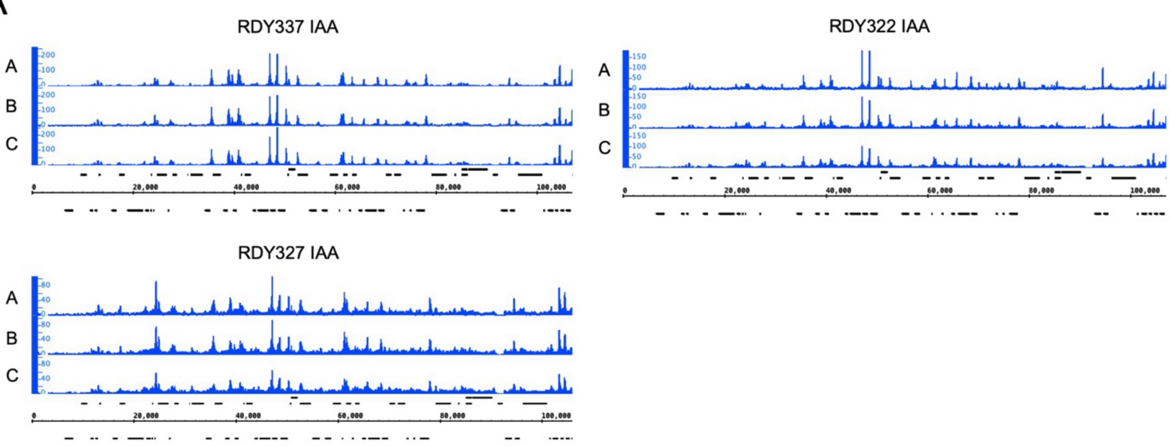
RDY337 IAA
RDY322 IAA
RDY327 IAA
A
B
C
20,000
40,000
60,000
80,000
100,000

B

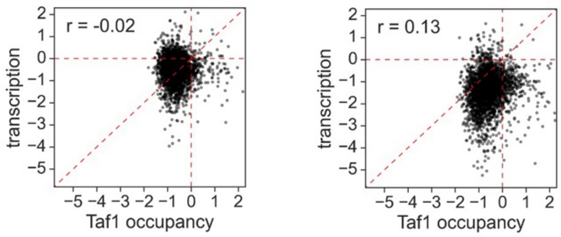
mBD1/2
mET(4)
r = -0.02
r = 0.13
transcription
Taf1 occupancy

**Fig. S7. The ET domain facilitates recruitment of TFIID to promoters.** (**A**) Comparison of ChEC-seq data for individual replicate experiments. Data for the first 100,000 bases of chromosome III is shown. (**B**) Comparison of changes in Taf1 promoter occupancy due to mutations in the BDs or the ET domain and corresponding changes in transcription. Pearson correlation coefficient (r) is shown.

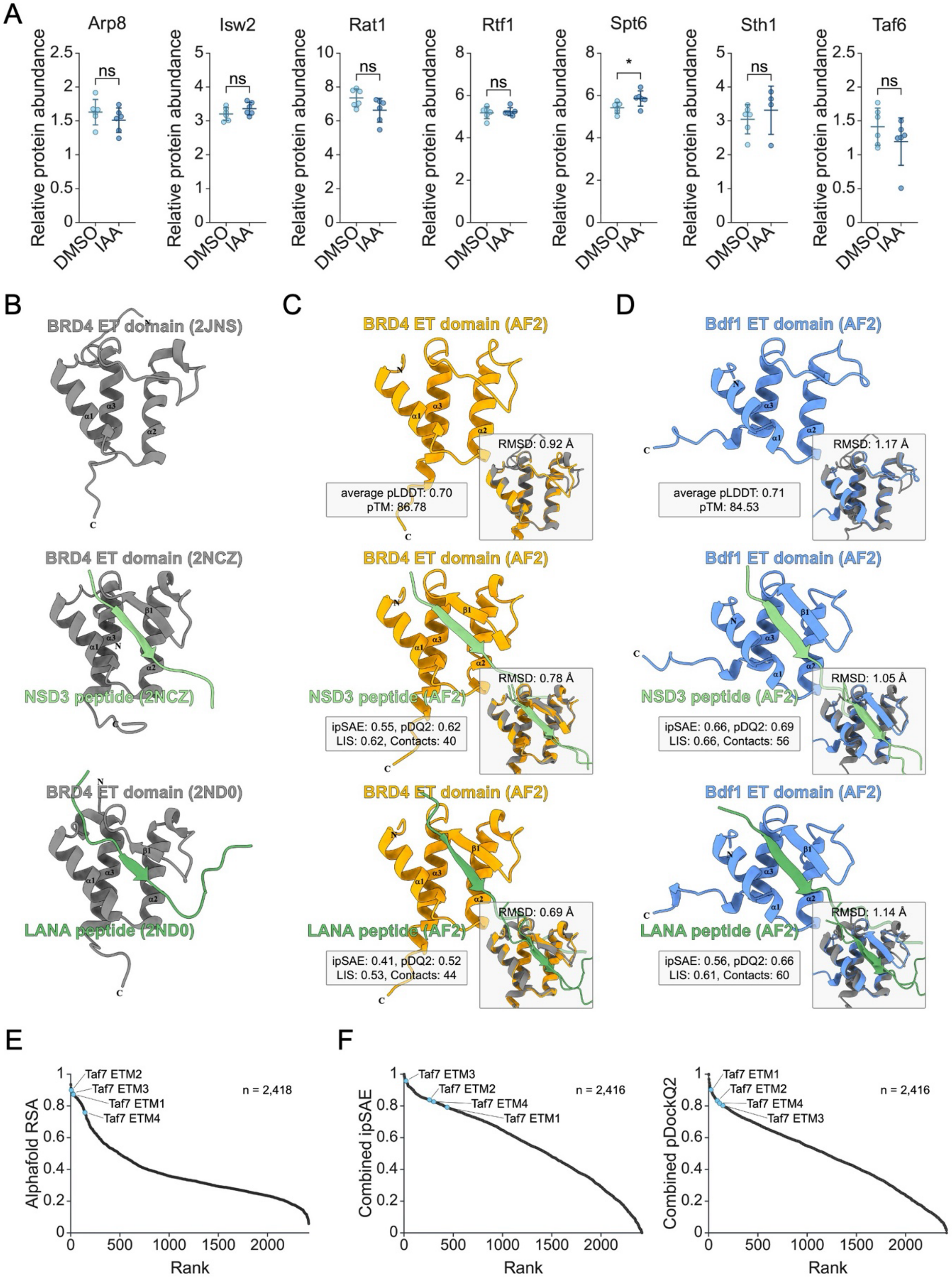
A
Arp8
Isw2
Rat1
Rtf1
Spt6
Sth1
Taf6
Relative protein abundance
DMSO
IAA
ns
*
B
BRD4 ET domain (2JNS)
BRD4 ET domain (2NCZ)
NSD3 peptide (2NCZ)
BRD4 ET domain (2ND0)
LANA peptide (2ND0)
C
BRD4 ET domain (AF2)
RMSD: 0.92 Å
average pLDDT: 0.70
pTM: 86.78
NSD3 peptide (AF2)
RMSD: 0.78 Å
ipSAE: 0.55, pDQ2: 0.62
LIS: 0.62, Contacts: 40
LANA peptide (AF2)
RMSD: 0.69 Å
ipSAE: 0.41, pDQ2: 0.52
LIS: 0.53, Contacts: 44
D
Bdf1 ET domain (AF2)
RMSD: 1.17 Å
average pLDDT: 0.71
pTM: 84.53
RMSD: 1.05 Å
ipSAE: 0.66, pDQ2: 0.69
LIS: 0.66, Contacts: 56
RMSD: 1.14 Å
ipSAE: 0.56, pDQ2: 0.66
LIS: 0.61, Contacts: 60
E
Alphafold RSA
Taf7 ETM2
Taf7 ETM3
Taf7 ETM1
Taf7 ETM4
n = 2,418
Rank
F
Combined ipSAE
n = 2,416
Combined pDockQ2

**Fig. S8. Protein levels are largely stable in whole cell extract following acute depletion of Bdf1/2, and structural predictions for the BRD4 or Bdf1 ET domains in association with ETMs are highly similar to experimental structures.** (**A**) Relative protein abundance in whole cell extract for selected proteins following Bdf1/2 depletion. Selected proteins are highlighted in Fig 5A. Taf2 and Rsc2 were not detected in whole cell extracts. Central lines indicate mean relative protein abundance. Error bars correspond to standard deviation of the mean ($n$ = 6). Results of a Welch's t-test are shown (* – $p$-value < 0.05, ns – not significant). (**B**) Previously determined experimental structures for the mouse BRD4 ET domain alone (PDB: 2JNS; (*81*)) (top panel) or the human BRD4 ET domain in association with characterized ETMs from NSD3 (PDB: 2NCZ (*24*)) (middle panel) or LANA (PDB: 2ND0 (*24*)) (bottom panel). (**C**) AlphaFold2 (AF2) predicted structures for the human BRD4 ET domain alone (top panel) or in association with ETMs from NSD3 (middle panel) or LANA (bottom panel). Confidence metrics (ipSAE, pDockQ2, and LIS) and the number of interfacial contacts supporting structural predictions are shown. Insets compare predicted and experimental structures with corresponding RMSD values. (**D**) AlphaFold2 (AF2) predicted structures for the Bdf1 ET domain alone (top panel) or in association with ETMs from NSD3 (middle panel) or LANA (bottom panel). Confidence metrics (ipSAE, pDockQ2, and LIS) and the number of interfacial contacts supporting the structural predictions are shown. Insets show comparisons between predicted and experimental structures and RMSD supporting the comparisons. (**E**) Scatter plot comparing Alphafold RSA scores and rank order for ETMs in nuclear proteins ($n$ = 2,418). Four putative ETMs in the Bdf1-interaction region of Taf7 are shown in blue. (**F**) Scatter plots comparing Combined ipSAE scores (left panel) or Combined pDockQ2 scores (right panel) and rank order for structural predictions of the Bdf1 ET domain in association with ETMs in nuclear proteins ($n$ = 2,416). Predicted structures involving four putative ETMs in the Bdf1-interaction region of Taf7 are shown in blue in both panels.

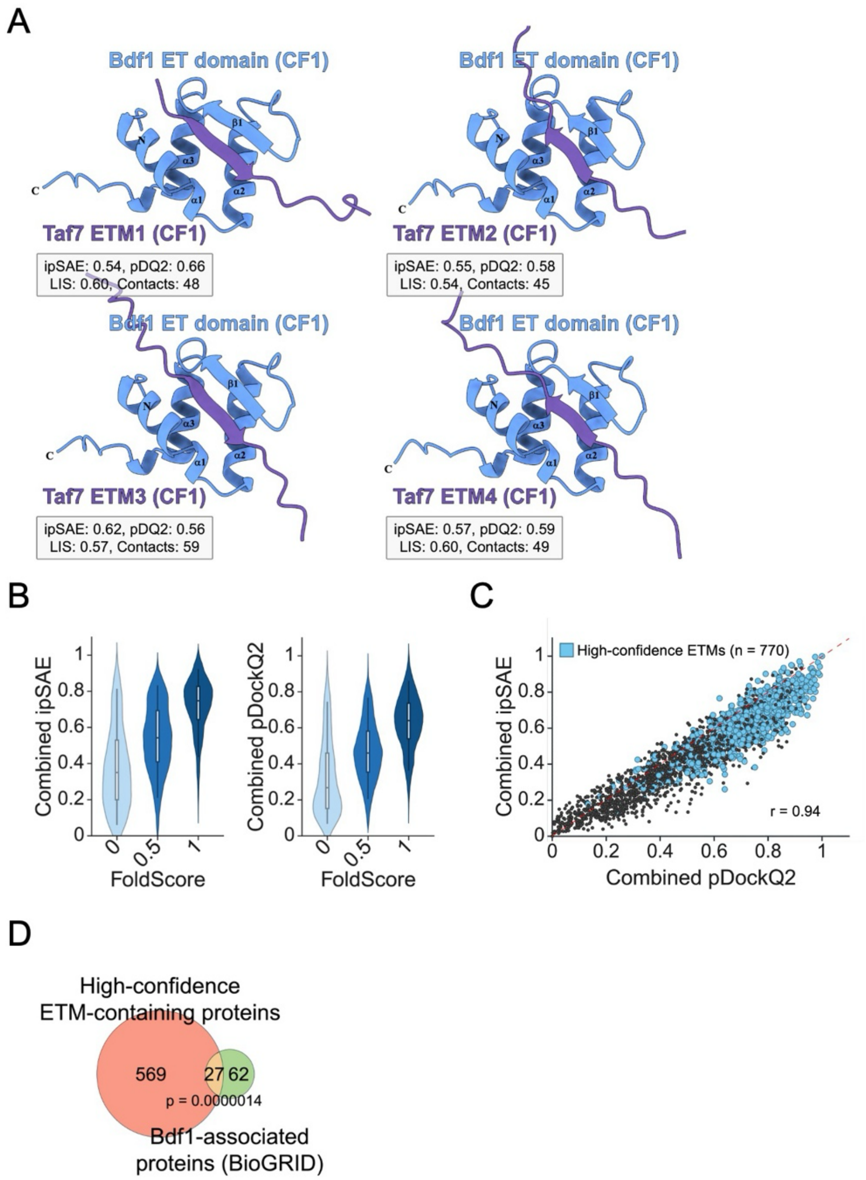
A
Bdf1 ET domain (CF1)
Taf7 ETM1 (CF1)
ipSAE: 0.54, pDQ2: 0.66
LIS: 0.60, Contacts: 48
Taf7 ETM2 (CF1)
ipSAE: 0.55, pDQ2: 0.58
LIS: 0.54, Contacts: 45
Taf7 ETM3 (CF1)
ipSAE: 0.62, pDQ2: 0.56
LIS: 0.57, Contacts: 59
Taf7 ETM4 (CF1)
ipSAE: 0.57, pDQ2: 0.59
LIS: 0.60, Contacts: 49
B
Combined ipSAE
Combined pDockQ2
FoldScore
C
High-confidence ETMs (n = 770)
r = 0.94
Combined pDockQ2
Combined ipSAE
D
High-confidence ETM-containing proteins
569
27
62
p = 0.0000014
Bdf1-associated proteins (BioGRID)

**Fig. S9. High-confidence ETMs are correlated with conditional interfacial β-sheet establishment, and proteins carrying these ETMs are enriched for functions related to transcription regulation and reported association with Bdf1.** (**A**) ColabFold (CF1) predicted structures for the Bdf1 ET domain in association with four putative ETMs in the Bdf1-interaction region of Taf7. Confidence metrics (ipSAE, pDockQ2, and LIS) and the number of interfacial contacts supporting structural predictions are shown. (**B**) Violin plots comparing Combined ipSAE scores (left panel) or Combined pDockQ2 scores (right panel) and FoldScore for structural predictions of the Bdf1 ET domain in association ETMs in nuclear proteins ($n$ = 2,416). (**C**) Scatter plot comparing Combined ipSAE scores and Combined pDockQ2 scores for structural predictions of the Bdf1 ET domain in association with ETMs in nuclear proteins ($n$ = 2,416). Structural predictions involving high-confidence ETMs ($n$ = 770) are shown in blue. High-confidence ETMs have an AlphaFold RSA $\geq$ 0.24 and a FoldScore = 1. Pearson correlation coefficient (r) is shown. (**D**) Bubble plot comparing selected, significantly enriched GO terms (Biological Process), fold enrichment of the GO term, and the number of genes associated with the GO term. Significance was calculated using a Fisher's exact test. Adjusted $p$-values were calculated using the Benjamini-Hochberg procedure. (**E**) Euler plot comparing high-confidence ETM-containing proteins ($n$ = 770) and proteins associated with Bdf1. The set of Bdf1-associated proteins was obtained from BioGRID (https://www.thebiogrid.org). Proteins with evidence for physical association were selected. Proteins identified in affinity capture RNA experiments or in a high-throughput Y2H study on coiled-coil domain interactions (*82*) were excluded. Results of a Fisher's exact test are shown.

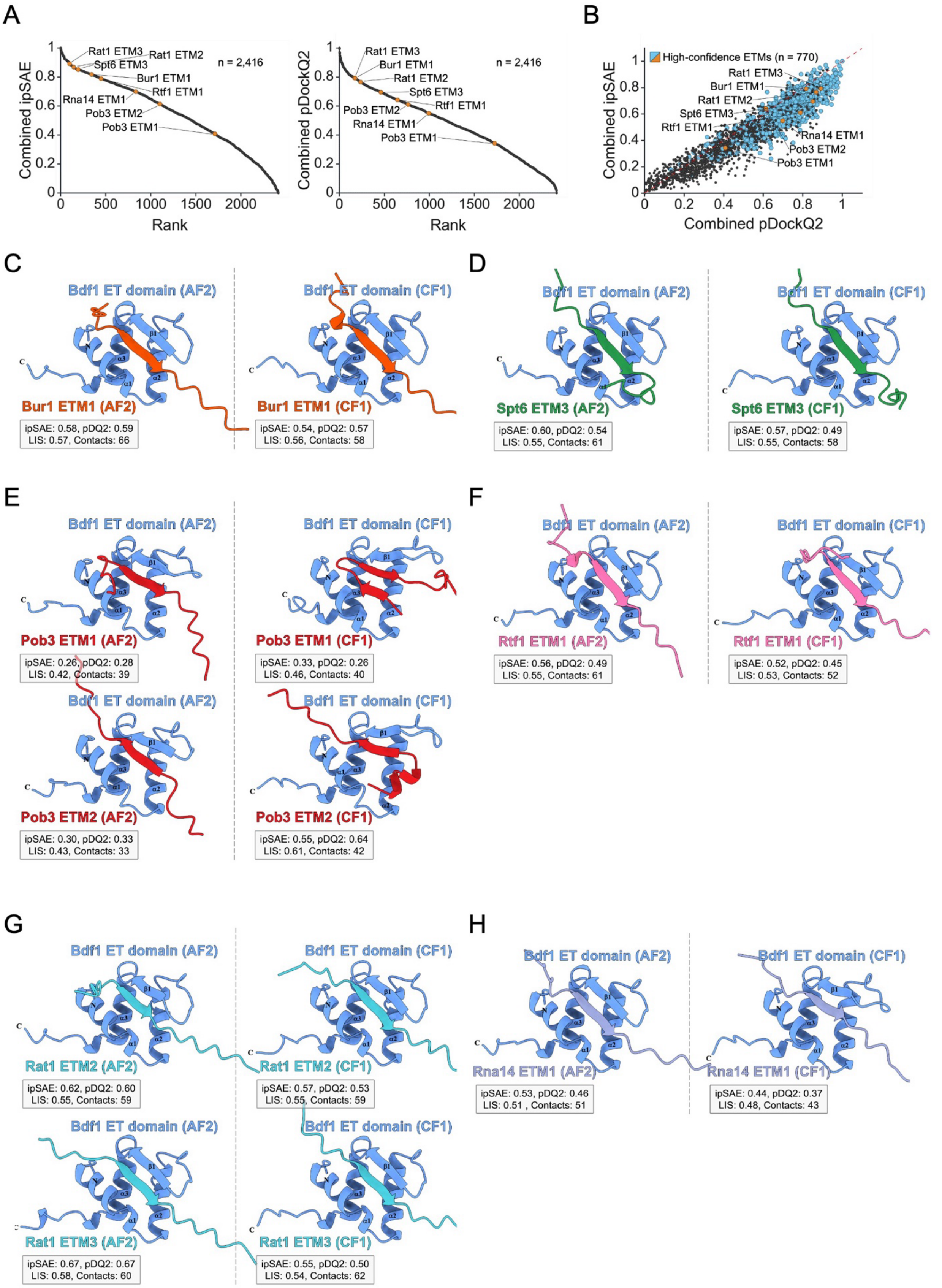
A
Combined ipSAE
Rank
n = 2,416
Rat1 ETM3
Rat1 ETM2
Spt6 ETM3
Bur1 ETM1
Rtf1 ETM1
Rna14 ETM1
Pob3 ETM2
Pob3 ETM1
Combined pDockQ2
B
High-confidence ETMs (n = 770)
Combined pDockQ2
C
Bdf1 ET domain (AF2)
Bdf1 ET domain (CF1)
Bur1 ETM1 (AF2)
Bur1 ETM1 (CF1)
ipSAE: 0.58, pDQ2: 0.59
LIS: 0.57, Contacts: 66
ipSAE: 0.54, pDQ2: 0.57
LIS: 0.56, Contacts: 58
D
Spt6 ETM3 (AF2)
Spt6 ETM3 (CF1)
ipSAE: 0.60, pDQ2: 0.54
LIS: 0.55, Contacts: 61
ipSAE: 0.57, pDQ2: 0.49
LIS: 0.55, Contacts: 58
E
Pob3 ETM1 (AF2)
Pob3 ETM1 (CF1)
ipSAE: 0.26, pDQ2: 0.28
LIS: 0.42, Contacts: 39
ipSAE: 0.33, pDQ2: 0.26
LIS: 0.46, Contacts: 40
Pob3 ETM2 (AF2)
Pob3 ETM2 (CF1)
ipSAE: 0.30, pDQ2: 0.33
LIS: 0.43, Contacts: 33
ipSAE: 0.55, pDQ2: 0.64
LIS: 0.61, Contacts: 42
F
Rtf1 ETM1 (AF2)
Rtf1 ETM1 (CF1)
ipSAE: 0.56, pDQ2: 0.49
LIS: 0.55, Contacts: 61
ipSAE: 0.52, pDQ2: 0.45
LIS: 0.53, Contacts: 52
G
Rat1 ETM2 (AF2)
Rat1 ETM2 (CF1)
ipSAE: 0.62, pDQ2: 0.60
LIS: 0.55, Contacts: 59
ipSAE: 0.57, pDQ2: 0.53
LIS: 0.55, Contacts: 59
Rat1 ETM3 (AF2)
Rat1 ETM3 (CF1)
ipSAE: 0.67, pDQ2: 0.67
LIS: 0.58, Contacts: 60
ipSAE: 0.55, pDQ2: 0.50
LIS: 0.54, Contacts: 62
H
Rna14 ETM1 (AF2)
Rna14 ETM1 (CF1)
ipSAE: 0.53, pDQ2: 0.46
LIS: 0.51 , Contacts: 51
ipSAE: 0.44, pDQ2: 0.37
LIS: 0.48, Contacts: 43

**Fig. S10. Structural modeling supports associations between the Bdf1 ET domain and key factors in transcription regulation.** (**A**) Scatter plots comparing Combined ipSAE scores (left panel) or Combined pDockQ2 scores (right panel) and rank order for predicted structures of the Bdf1 ET domain in association with ETMs in nuclear proteins ($n$ = 2,416). Predicted structures involving high-confidence ETMs in Bur1, Spt6, Pob3, Rtf1, Rat1, or Rna14 are shown in orange in both panels ($n$ = 8). (**B**) Scatter plot comparing Combined ipSAE scores and Combined pDockQ2 scores for structural predictions of the Bdf1 ET domain in association with ETMs in nuclear proteins ($n$ = 2,416). Structural predictions involving high-confidence ETMs in Bur1, Spt6, Pob3, Rtf1, Rat1, or Rna14 are shown in orange ($n$ = 8). Structural predictions involving the remaining high-confidence ETMs are shown in blue ($n$ = 762). (**C-H**) Predicted structures for the Bdf1 ET domain in association with high-confidence ETMs in Bur1, Spt6, Pob3, Rtf1, Rat1, or Rna14. AlphaFold2 (AF2) and ColabFold (CF1) predicted structures are shown in right and left panels, respectively. Confidence metrics (ipSAE, pDockQ2, and LIS) and the number of interfacial contacts supporting structural predictions are shown.

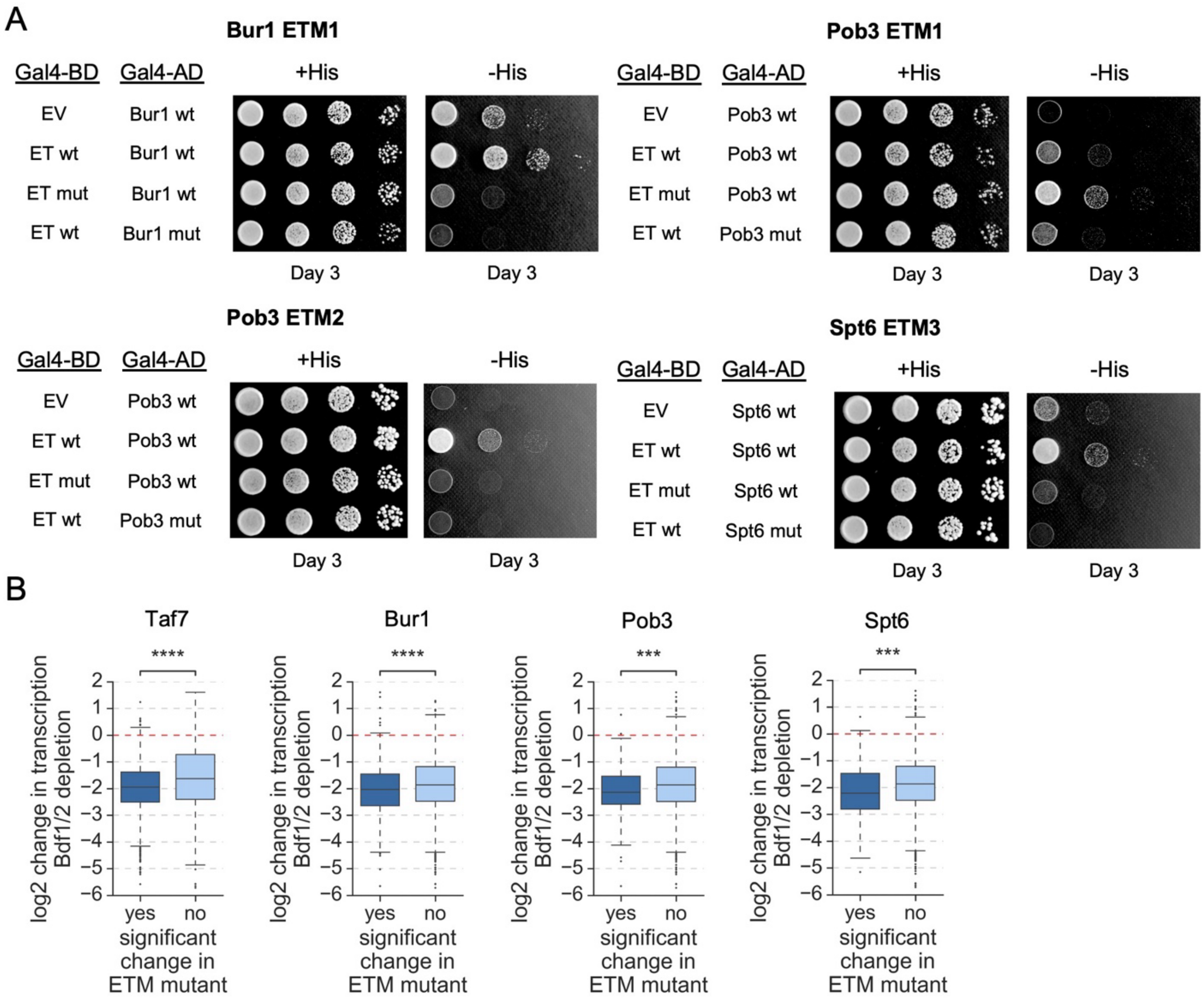

A
Bur1 ETM1
Gal4-BD
Gal4-AD
+His
-His
EV
Bur1 wt
ET wt
Bur1 wt
ET mut
Bur1 wt
ET wt
Bur1 mut
Day 3
Day 3
Pob3 ETM1
Gal4-BD
Gal4-AD
+His
-His
EV
Pob3 wt
ET wt
Pob3 wt
ET mut
Pob3 wt
ET wt
Pob3 mut
Day 3
Day 3
Pob3 ETM2
Gal4-BD
Gal4-AD
+His
-His
EV
Pob3 wt
ET wt
Pob3 wt
ET mut
Pob3 wt
ET wt
Pob3 mut
Day 3
Day 3
Spt6 ETM3
Gal4-BD
Gal4-AD
+His
-His
EV
Spt6 wt
ET wt
Spt6 wt
ET mut
Spt6 wt
ET wt
Spt6 mut
Day 3
Day 3
B
Taf7
Bur1
Pob3
Spt6
****
****
***
***
log2 change in transcription Bdf1/2 depletion
2
1
0
−1
−2
−3
−4
−5
−6
yes
no
significant change in ETM mutant

**Fig. S11. The loss of interactions between the Bdf1 ET domain and ETMs in Bur1, Pob3, and Spt6 contributes to the overall defects observed with Bdf1/2 depletion.** (**A**) Y2H analysis of interactions between the Bdf1 ET domain and high-confidence ETMs in Bur1, Pob3, and Spt6. Plasmids expressing the Bdf1 ET domain (WT or mut), fragments of Bur1, Pob3, or Spt6 that cover the indicated ETMs (WT or mut), or an empty vector (EV) were co-transformed into the host strain. Cells were serially diluted, spotted on the indicated media, and imaged after three days of growth ($n = 3$). ET mut – ET domain with four conserved acidic residues substituted with alanine. Basic residues in all ETM mutants are substituted with alanine. Amino acid coordinates of Bur1, Pob3, and Spt6 peptides used are as follows: Bur1 ETM1 – 2-62, Pob3 ETM1 – 239-383, Pob3 ETM2 – 381-552, Spt6 ETM3 – 1249-1451. (**B**) Log2 changes in transcription following depletion of Bdf1/2 for 25 min. Genes are categorized into two groups based on the statistical significance of the changes observed in relation to ETM mutations in the indicated factors (Taf7, Bur1, Pob3, or Spt6). Results of a Welch's t-test are shown (*** – $p$-value < 0.001, **** – $p$-value < 0.0001).

**Supplementary tables associated with this study:**

**Table S1. List of plasmids used in this study.**

**Table S2. List of strains used in this study.**

**Table S3. Results of SLAM-seq experiments (Figs. 2A-F, 6C, S2, S3, S4A-B, S4D-E and S11B).**

**Table S4. Log2 changes in Bdf1 promoter occupancy due to *BDF1* mutations (Figs. 3 and S6).**

**Table S5. Results of chromatin mass spectrometry experiments (Figs. 5A and S8A).**

**Table S6. Summary of ET-interacting motif (ETM) discovery (Fig. 5B).** A list of 2418 ETMs identified in yeast nuclear proteome is shown in tab D.

**Table S7. Summary of structural predictions and associated analysis (Figs. 5D, S8B-F, S9 and S10).** Lists of high-confidence ETMs and associated proteins are shown in tabs E and F, respectively.
